# Oxidative stress pathogenically remodels the cardiac myocyte cytoskeleton via structural alterations to the microtubule lattice

**DOI:** 10.1101/2020.06.30.174532

**Authors:** Rebecca R. Goldblum, Mark McClellan, Cody Hou, Brian R. Thompson, Kyle White, Hluechy X. Vang, Houda Cohen, Joseph M. Metzger, Melissa K. Gardner

## Abstract

In the failing heart, the cardiac myocyte microtubule network is remodeled, which increases cellular stiffness and disrupts contractility, contributing to heart failure and death. However, the origins of this deleterious cytoskeletal reorganization are unknown. We now find that oxidative stress, a condition characteristic of failing heart cells, leads to cysteine oxidation of microtubules. Further, our electron and fluorescence microscopy experiments revealed regions of structural damage within the oxidized microtubule lattice. These damaged regions led to the lengthening, realignment, and acetylation of dynamic microtubules within cardiac myocytes. Thus, we found that oxidative stress acts inside of cardiac myocytes to facilitate a dramatic, pathogenic shift from a dynamic, multifaceted microtubule network into a highly acetylated, longitudinally aligned, and static microtubule network. Our results demonstrate how a disease condition characterized by oxidative stress can trigger a molecular oxidation event, which propagates a toxic cellular-scale transformation of the cardiac myocyte microtubule network.

## Introduction

Oxidative stress is a hallmark of many cardiac pathologies, including the prolonged ischemia and reperfusion associated with ischemic heart disease, cardiac hypertrophy, and heart failure (Hori and Nishida, 2009; Slezak et al., 1995; Thompson and Hess, 1986). Global ischemia in the heart leads to a ∼250% increase in reactive oxygen species, suggesting that oxidative stress is an important driving force leading to changes in the cellular cytoskeleton that accompany ventricular remodeling in ischemic heart disease and heart failure (Hein et al., 2000; Slezak et al., 1995; Wang et al., 2008). Similarly, oxidative-stress exposed cells isolated from healthy tissue undergo reorganization and/or gradual coarsening of the microtubule network (Drum et al., 2016; Valen et al., 1999; Wei-Guo and Qi-Ping, 2014). Alterations in the microtubule cytoskeletal network suggest that oxidative stress can directly modulate microtubule dynamics or stability (Drum et al., 2016).

Importantly, pathological changes in microtubule cytoskeletal organization have been shown to increase cellular stiffness and lead to contractile dysfunction of cardiac myocytes (Bauer et al., 2000; Cooper, 2006; Hein et al., 2000; Howarth et al., 1999; Iwai et al., 1990; Nishimura et al., 2006; Sato et al., 1997; Tagawa et al., 1998; Tsutsui et al., 1993, 1994; Webster, 2002; White, 2011). Furthermore, microtubule reorganization hinders intracellular transport of proteins to their targets at the cell surface (Drum et al., 2016; Prins et al., 2016). Many of these proteins, such as the potassium channels, Kv4.2, Kv4.3, and Kv1.5, are involved in cardiac myocyte electrical signaling. Failure of these channels to reach the plasma membrane leads to heart arrhythmia, also known as an irregular heartbeat (Drum et al., 2016; Prins et al., 2016; Zadeh et al., 2009). While it has been shown that oxidative stress alters microtubule networks, promotes increased cardiac myocyte stiffness, and obstructs microtubule-dependent transport, the nature and mechanism of oxidative stress-mediated remodeling of the microtubule cytoskeleton is poorly understood.

Microtubules are cylindrical polymers of α/β-tubulin heterodimers that form a complex cytoskeletal network throughout the cytoplasm (Desai and Mitchison, 1997; Kirschner and Mitchison, 1986, 2002; Mitchison and Kirschner, 1984; Ohi and Zanic, 2016). Microtubule plus-end dynamics are biphasic, characterized by periods of slow microtubule growth (polymerization) followed by intervals of abrupt shortening (depolymerization). Microtubule length and function depends on the tight regulation of microtubule growth and shortening rates, as well as the frequencies of “catastrophe” events – the transition from growth to shortening, and “rescue” events – the transition from shortening to growth.

During microtubule growth, GTP-bound tubulin subunits are added to microtubule plus-ends (Nogales et al., 1998). Once these GTP-tubulin subunits are buried within the microtubule lattice, they are hydrolyzed to GDP-tubulin (Desai and Mitchison, 1997; Howard and Hyman, 2003; Mitchison and Kirschner, 1984). This results in a GTP-tubulin ‘cap’ at the growing end of a predominantly GDP-tubulin microtubule filament. Importantly, GTP-tubulin is more stable within the microtubule lattice than GDP-tubulin, and thus the GTP-tubulin cap stabilizes microtubules and prevents rapid depolymerization (Desai and Mitchison, 1997). Recently, it has been shown that patches of GTP tubulin that are present along the length of the GDP-tubulin microtubule lattice may increase the likelihood of rescue events, thus promoting net microtubule elongation (Aumeier et al., 2016; Dimitrov et al., 2008; de Forges et al., 2016; Gardner, 2016; Tropini et al., 2012). In cells, microtubules can be post-translationally acetylated, which increases their resistance to breakage, and thus also increases their longevity (Akella et al., 2010; Eshun-Wilson et al., 2019; Kormendi et al., 2012; Maruta et al., 1986; Portran et al., 2017; Shida et al., 2010; Xu et al., 2017).

In this work, we found that oxidative stress acts to facilitate a dramatic and maladaptive longitudinal realignment and acetylation of microtubules inside of cardiac myocytes. Specifically, we found that microtubules subjected to oxidative stress undergo cysteine oxidation. Our electron and fluorescence microscopy experiments revealed that the oxidized microtubules had structural damage within the cylindrical tubulin lattice, consisting of holes and lattice openings. For dynamic microtubules, incorporation of stabilizing GTP-tubulin into these damaged lattice regions led to an increased frequency of rescue events (the transition from shortening to growth), and thus longer microtubules. Within cardiac myocytes, this lengthening of dynamic microtubules allows for their interaction with regularly spaced Z-disks, leading to longitudinal alignment of microtubules within the cardiac myocyte sarcomeric structure. Further, the damaged lattice regions generated via oxidative stress increased the efficiency of microtubule acetylation: we found that the longitudinally aligned, oxidized microtubules were highly acetylated, which increases microtubule longevity and thus further reinforces deleterious microtubule network remodeling. Thus, oxidative stress triggers a dramatic, pathogenic shift from a dynamic, multifaceted microtubule network, into a highly acetylated, longitudinally aligned, and static microtubule network inside of cardiac myocytes, likely contributing to increased cellular stiffness and contractile dysfunction. Our results provide insight into myocardial changes in Ischemic Heart Disease by describing a mechanism for the dramatic remodeling of the microtubule cytoskeletal network within cardiac myocytes subjected to oxidative stress.

## Results

### Oxidative stress increases longitudinal alignment of the microtubule network in cardiac myocytes

To examine microtubule organization in cardiac myocytes subjected to oxidative stress, rat ventricular cardiac myocytes were isolated and adhered to coverslips placed in 6-well plates, as previously described (Thompson et al., 2019). We then subjected isolated rat cardiac ventricular myocytes to 100-1000 µM hydrogen peroxide (H_2_O_2_), a reactive oxygen species that is upregulated in ischemic cardiac myocytes (Slezak et al., 1995). The physiological intracellular H_2_O_2_ concentration has been estimated to be 0.1-15 µM (Fernández-Puente et al., 2020; Giorgio et al., 2007; Sies, 2017; Slezak et al., 1995), but can be as high as 50 µM in the context of cardiac ischemia and reperfusion (Slezak et al., 1995). However, the amount of H_2_O_2_ that actually enters the cell from H_2_O_2_ in surrounding media has been reported to be 100 to 650-fold lower than the concentration in the media (Huang and Sikes, 2014; Sies, 2017). Therefore, we predicted that media H_2_O_2_ concentrations of 100-1000 µM would lead to low µM internal cellular H_2_O_2_ concentrations, a range that should approximate pathophysiological conditions in cardiac ischemia. Importantly, this range of media H_2_O_2_ concentrations has been widely used to evaluate the effects of oxidative stress in cells (Kratzer et al., 2012; Valen et al., 1999; Wei-Guo and Qi-Ping, 2014; Xie et al., 2014; Zhang et al., 2018; Zhao et al., 2019).

Thus, cardiac myocytes were treated for 1 hour in M199 media alone, or in M199 media containing 0–1 mM H_2_O_2_. To characterize the microtubule cytoskeletal network, cardiac myocytes were then fixed in 1% PFA, stained with anti-α-tubulin antibody, and imaged with confocal microscopy. Control cardiac myocytes (Fig. 1A, left) displayed a sparse, complex network of microtubules, similar to previous reports (Robison and Prosser, 2017). In contrast, cardiac myocytes treated with 100 μM H_2_O_2_ (Fig. 1A, center) and 1 mM H_2_O_2_ (Fig. 1A, right) had a dramatic cytoskeletal reorganization: microtubules were uniformly arranged such that the microtubules were lengthened and longitudinally reoriented along the long axis of the cardiac myocyte. This result is similar to the pathological microtubule remodeling reported in failing cardiac myocytes (Tagawa et al., 1998; Tsutsui et al., 1993, 1994).

**Figure 1:**
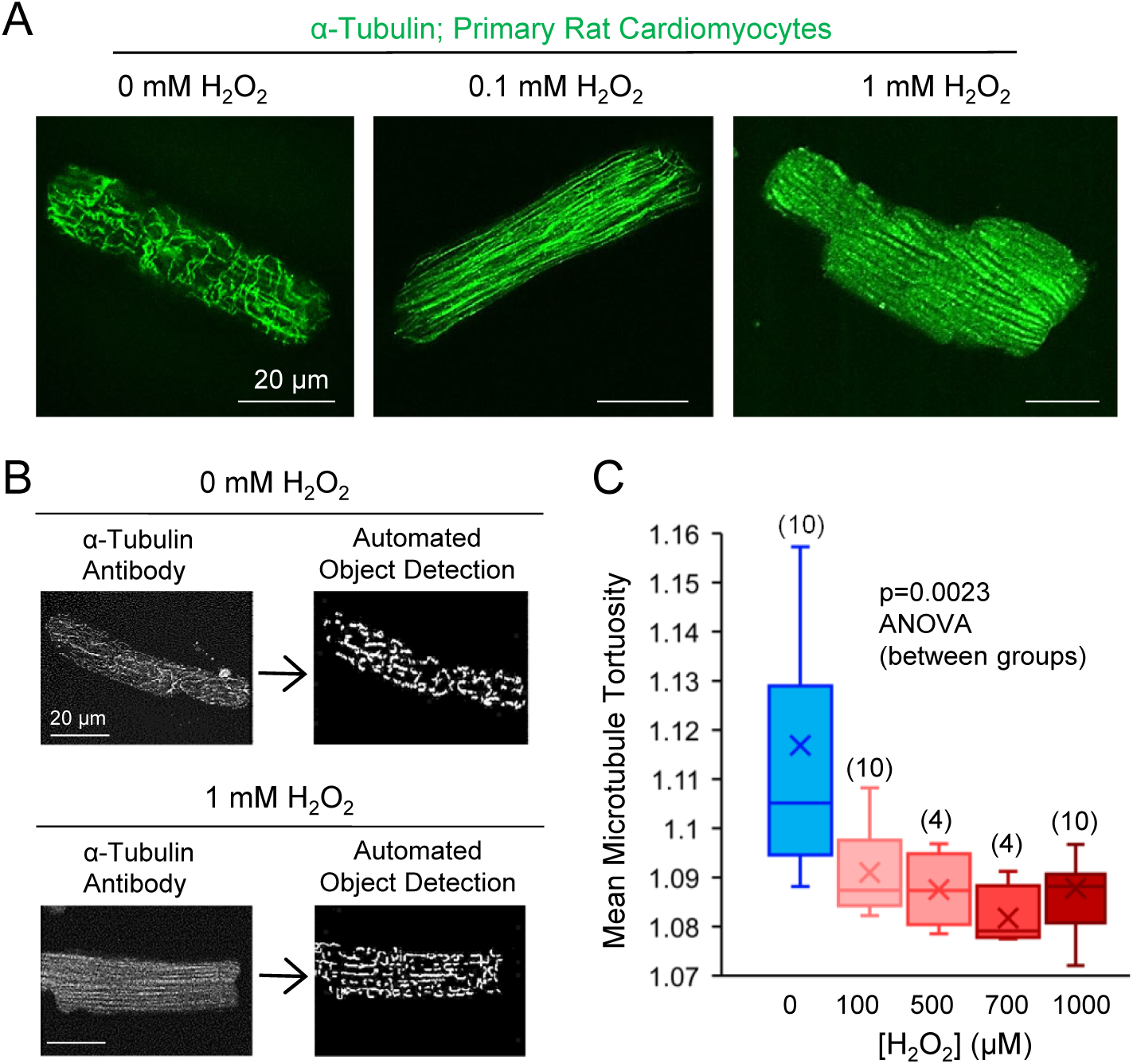
Oxidative stress leads to longitudinal alignment of the microtubule cytoskeleton in cardiac myocytes. (A) Cardiac myocytes exposed to 0 mM, 0.1 mM, or 1 mM H_2_O_2_ for 1h before fixation in 1% PFA and immunostaining for α-tubulin (green). (B) Tortuosity is measured by automatically detecting microtubules within cardiac myocytes. (C) Mean microtubule tortuosity is reduced in cardiac myocytes with H_2_O_2_ exposure, consistent with the straightening and longitudinal alignment of microtubules. Crosses: Mean; Bar: Median; Box: first-third quartiles. Sample sizes as shown in parenthesis, numbers represent number of cells analyzed.

To quantify this observation, we estimated the “tortuosity” of microtubules within each treatment group. Tortuosity of a microtubule can be defined as:

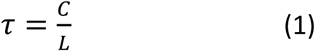

where *τ* represents tortuosity, *C* represents the total path length of a curved microtubule, and *L* represents the distance between the two ends of the microtubule. Thus, a straight line will have a minimal tortuosity of 1, while a circle will have a tortuosity of infinity. Importantly, a decreased tortuosity value indicates that microtubules are straighter and more aligned. By using automated object detection to detect microtubules within an image (Fig. 1B), the tortuosity was quantified for each microtubule segment, and the average microtubule tortuosity was computed across all of the segments within each cardiac myocyte. The mean microtubule tortuosity per cardiac myocyte was then compared for different treatment groups (Fig. 1C). We found that the tortuosity of microtubule segments was dramatically decreased at even the lowest H_2_O_2_ concentration tested (Fig. 1B, right. p=0.0023 ANOVA, difference between groups, sample sizes=number of cells analyzed). This decrease in tortuosity suggests that oxidative stress leads to longitudinal reorientation and lengthening along the long axis of the cardiac myocyte, a pathological condition that could disrupt cellular contractility.

### Oxidative stress directly increases microtubule rescue frequency in cell-free experiments

To investigate the intrinsic mechanism for this longitudinal microtubule alignment, we turned to cell-free reconstitution experiments. Because it is likely that growing microtubule ends interact with cardiac myocyte intermediate filaments to alter their dynamics, we reasoned that cell-free reconstitution experiments would allow us to evaluate the direct effect of oxidative stress on microtubule dynamics, at single-microtubule, nanoscale resolution, and independently of cardiac myocyte health or the interactions of growing microtubule ends with cardiac myocyte sarcomeric structures.

Thus, tubulin was purified from swine brain, and microtubule-associated proteins were removed through two cycles of polymerizations in high-salt buffer (Castoldi and Popov, 2003).The purified tubulin was then labeled with Alexa-488, and green Alexa-488 microtubules were imaged growing from stabilized, coverslip-attached rhodamine-labeled (red) microtubule “seeds” using Total Internal Reflection Fluorescence (TIRF) microscopy (Gell et al., 2010) (Fig. 1A, top; Supplementary Movie S1; see materials and methods).

Movies were collected at 0.2 frames/s over 30-60 min intervals (Supplementary Movie S1), and then converted into “kymographs” to analyze the growth and shortening behavior of the microtubules (Fig. 1A, bottom). Thus, the growth and shortening rates of dynamic microtubules were estimated, along with the frequency of switching between these two states. To determine whether oxidative stress would directly alter microtubule dynamics, these parameters were then quantified for growing microtubules that were exposed to increasing concentrations of H_2_O_2_ (Fig. 1A, bottom). We found that microtubule growth and shortening rates were not significantly altered by increasing H_2_O_2_ concentrations (growth: Fig. 2B, p=0.43, R^2^=0.213; shortening: Fig. 2C, p=0.57, R^2^=0.18), similar to a previous study in cardiac myocytes (Drum et al., 2016). Thus, oxidative stress does not alter the kinetics of microtubule assembly and disassembly.

**Figure 2:**
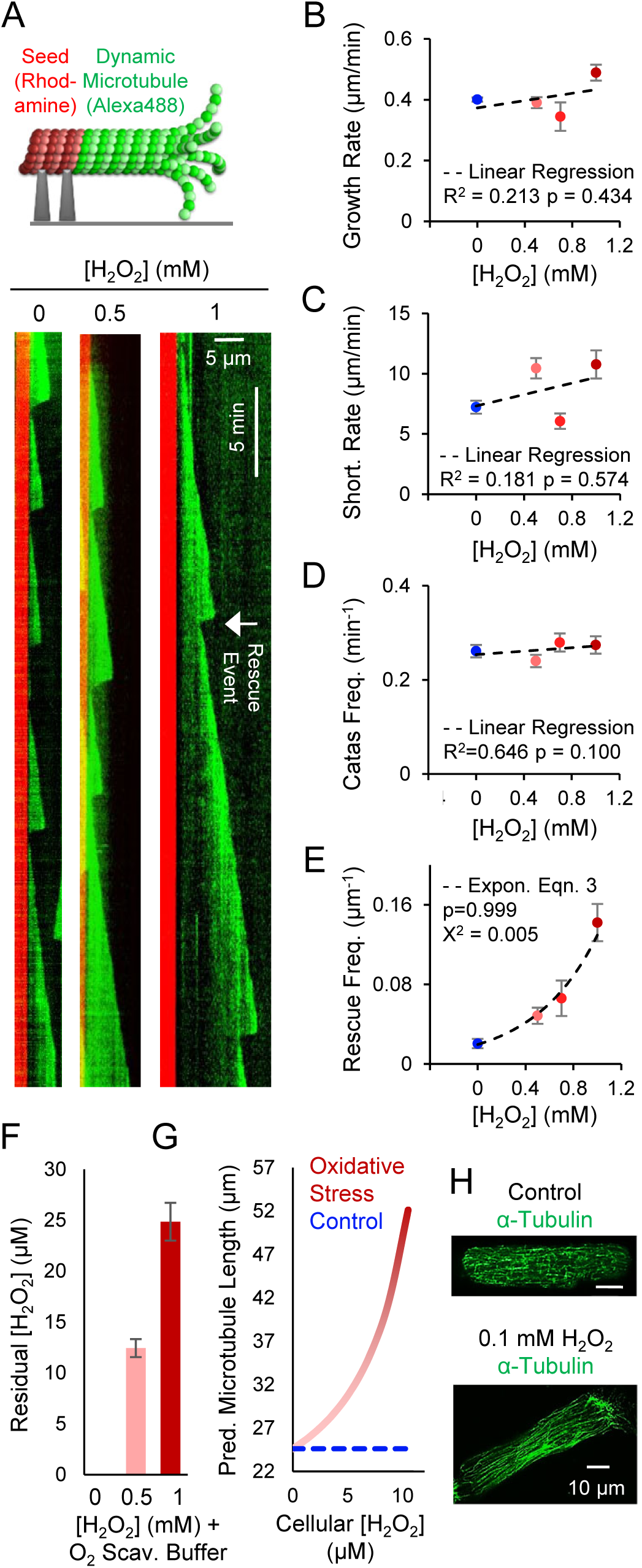
Oxidative stress increases dynamic microtubule length by increasing rescue frequency. (A) (Top) Schematic of TIRF experiments to measure microtubule length dynamics. (Bottom) Representative kymographs of dynamic microtubules in the presence of 0 mM H_2_O_2_ (left), 0.5 mM H_2_O_2_ (middle), and 1 mM H_2_O_2_ (right) (red, GMPCPP-stabilized microtubule seeds; green, Alexa-Fluor-488 dynamic microtubule extensions; white arrow: rescue event). (B) Microtubule growth rate as a function of H_2_O_2_ concentration (n=280, 265, 149, 123 from 0 to 1 mM on graph). (C) Microtubule shortening rate as a function of H_2_O_2_ concentration (n=41, 39, 41, 39 from 0 to 1 mM on graph). (D) Microtubule catastrophe frequency as a function of H_2_O_2_ concentration (n=280, 265, 149, 123 from 0 to 1 mM on graph). (E) Microtubule rescue frequency, calculated as the frequency of a rescue event per catastrophe, and normalized to mean microtubule length at catastrophe, n=280, 265, 149, 123 catastrophe events from 0 to 1 mM on graph, absolute number of rescue events observed, from 0 to 1 mM on graph: n=20, 45, 26, 90. (F) Measurements of residual H_2_O_2_ concentration in the oxygen scavenging imaging buffer used in panels A-E. Average shown based on three independent repeats with Amplex Red assay. (G) Predicted microtubule length as a function of increasing oxidative stress in cardiac myocytes, based on Eqns. 2,3, and panel F. (G) Typical images of tubulin staining in cardiac myocytes appear consistent with increased microtubule length predictions. All error bars, in all panels, mean ± S.E.M.

We then measured the growth lifetime of each microtubule prior to shortening, and inverted these lifetime values to determine the “catastrophe frequency” (min^−1^). We found that the catastrophe frequency of microtubules was unchanged with increasing H_2_O_2_ concentrations (Fig. 1D, p=0.89, R^2^=0.65), indicating that oxidative stress does not alter average microtubule growth lifetime. A previous study in cardiac myocytes reported an increase in catastrophe frequency for microtubule subjected to oxidative stress (Drum et al., 2016), but this difference could be due in part to the use of EB3 overexpression to visualize microtubule dynamics in cardiac myocytes, as EB proteins themselves increase the catastrophe frequency of dynamic microtubules (Zanic et al. 2013), and so their overexpression could alter sensitivity of catastrophe frequency to oxidative stress.

Finally, we measured the frequency of “rescue” events, in which a post-catastrophe microtubule would cease shortening and begin to grow again, prior to complete depolymerization to the red stabilized “seed” (Fig. 1A, bottom-right, white arrow)). The rescue frequency per catastrophe was normalized to the maximum microtubule length prior to catastrophe, and then plotted as a function of increasing H_2_O_2_ concentration (Fig. 2E). Strikingly, we found that rescue frequency increased exponentially with increasing H_2_O_2_ concentrations (Fig. 2E, Chi-squared goodness of fit test to exponential curve, p=1, χ^2^ test statistic=0.005), with a ∼7-fold increase in rescue frequency at 1 mM H_2_O_2_. This trend is consistent with a previous study in cardiac myocytes (Drum et al., 2016).

### Increase in microtubule rescue frequency with oxidative stress predicts longer microtubules in cardiac myocytes

To estimate the effect of increased rescue frequency on microtubule length in cardiac myocytes, we applied the Dogterom and Leibler equation (Leibler and Dogterom, 1993), which has been widely used to estimate the average length of microtubules based on the measured parameters of microtubule dynamics, as follows:

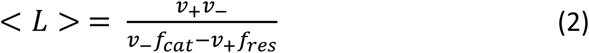

where *v*_+_ is the growth rate, *v*_*-*_ is the shortening rate, *f*_*cat*_ is the rescue frequency, and *f*_*res*_ is the rescue frequency. Using Eqn. 1 and the previously measured basal parameters in untreated mouse ventricular myocytes (Drum et al., 2016), we predicted the average microtubule length in untreated cardiac myocytes as ∼25 μm (Fig. 2F, blue), which appears roughly consistent with our rat cardiac myocyte images in control cells (Fig. 2G, top). We then predicted the effect of H_2_O_2_ exposure on the dynamic microtubule length in cardiac myocytes using our experimentally determined relationship between H_2_O_2_ concentration and rescue frequency, as follows (Fig. 2E):

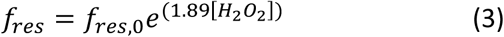

where *f*_*res*_ is the rescue frequency in the presence of H_2_O_2_, and *f*_*res,0*_ is the previously measured cardiac myocyte microtubule rescue frequency in the absence of H_2_O_2_ (Drum et al., 2016). However, because the experiments in Fig. 2B-E were performed in the presence of an oxygen scavenging buffer, we first used a previously published Amplex Red assay to estimate the residual H_2_O_2_ concentration in these assays (Schlieve et al., 2006). Similar to our estimates of internal cellular H_2_O_2_ concentration relative to media H_2_O_2_ concentration, the residual H_2_O_2_ concentrations are low μM in the cell-free assay that includes oxygen scavenging buffer (Fig. 2F). Thus by substituting Eqn. 3 into Eqn. 2, we predicted the relationship between cellular H_2_O_2_ concentration and microtubule lengths in cardiac myocytes (Fig. 2G, red). Similar to the effect on rescue frequency (Fig. 2E), Eqn. 3 predicts that, if allowed to grow without obstruction or interaction with cellular structures, dynamic microtubule lengths in cardiac myocytes would increase exponentially in response to increasing concentrations of reactive oxygen species in the cell, similar to our qualitative observations of the microtubule network in H_2_O_2_-treated cardiac myocytes (Fig. 2H, bottom).

### Oxidative stress leads to cysteine oxidation of microtubules

We then sought to elucidate a mechanism for how oxidative stress markedly increases microtubule rescue frequency, and thus increases the predicted dynamic microtubule length in cardiac myocytes. Tubulin dimers contain 20 cysteine residues and the oxidation of several have been shown to alter microtubule polymerization (Clark et al., 2014; Landino et al., 2002, 2011). We therefore tested whether H_2_O_2_ exposure oxidizes microtubules. Purified microtubules were grown in the presence of DCP-Rho1, which selectively binds sulfenic acid (Poole et al., 2007), and is thus a positive indicator of cysteine oxidation. We found that microtubules grown in the presence of 0.5 mM H_2_O_2_ revealed bright green fluorescent DCP-Rho1 binding (Fig. 3A, left) which was near non-detectable in the H_2_O_2_-free controls (Fig. 3A, Left). By using our previously published MATLAB code to quantify the fraction of microtubule area occupied by fluorescent DCP-Rho1 (Coombes et al., 2016; Reid et al., 2017), we found that DCP-Rho1 binding was increased by 16-fold with H_2_O_2_ exposure (Fig. 3A, right; see materials and methods). Thus, oxidative stress leads to cysteine oxidation of microtubules.

**Figure 3:**
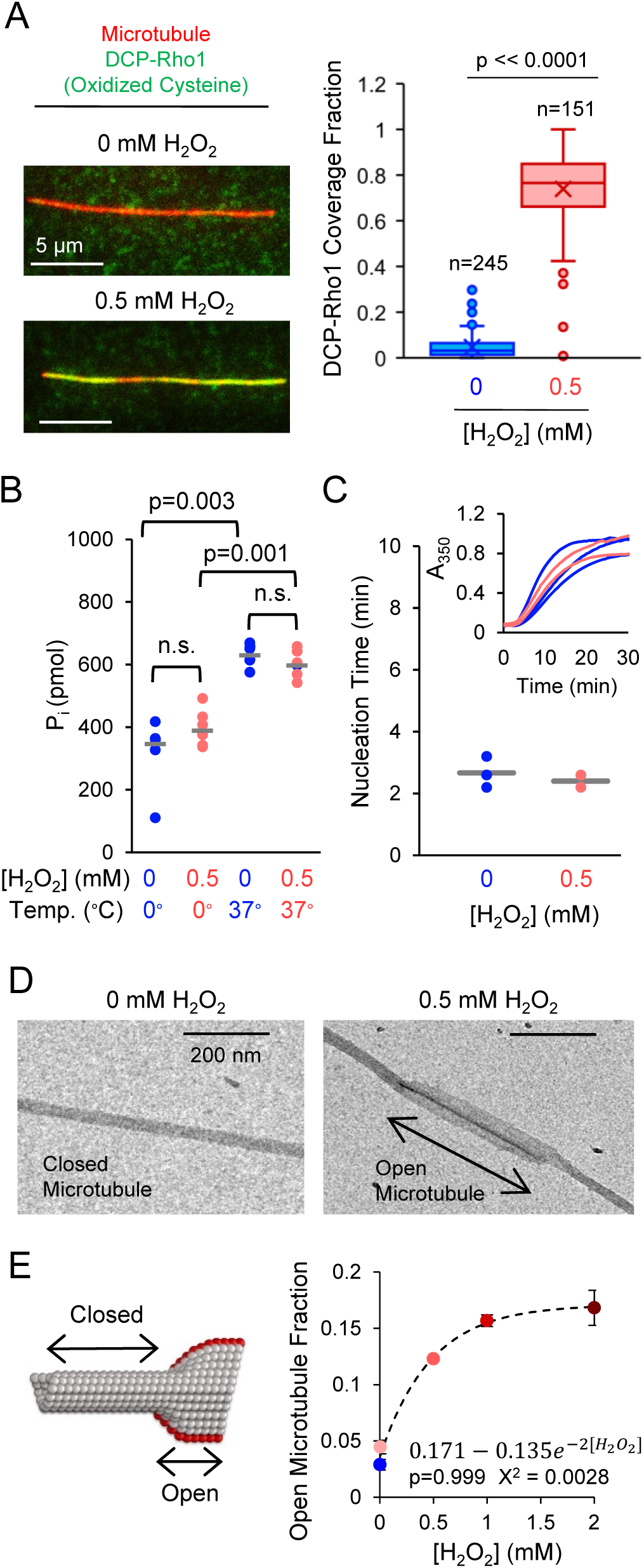
H_2_O_2_ exposure leads to cysteine oxidation and to microtubule damage. (A) Right: TIRF microscopy of GDP-tubulin microtubules (red) grown in the presence of DCP-Rho1 (green), an indicator of cysteine oxidation, in 0 mM H_2_O_2_ (top) or 1 mM H_2_O_2_ (bottom). Left: Quantification of DCP-Rho1 coverage area fraction on microtubules. Crosses: Mean; Bar: Median; Box: first-third quartiles; sample sizes represent number of images, many microtubules per image (B) GTPase assay. Dot plot showing the molar concentration of inorganic phosphate released over 2.5 h in 0 or 37°C, and in 0 or 0.5 mM H_2_O_2_. Sample sizes from left to right on graph: n= 6, 7, 7, 6, each number represents an independent experiment. Horizontal bars: mean; p-values calculated using Mann-Whitney Test. (C) Nucleation time for control (blue) and 0.5 mM H_2_O_2_-treated (red) microtubules. Inset: Turbidity, or absorbance at 350 nm, to monitor polymer formation over 30 min. Horizontal bars: mean. (D) Representative electron microscopy images of GDP-tubulin microtubules treated with 0 mM H_2_O_2_ (left) and 0.5 mM H_2_O_2_ (right). Black arrow, opened microtubule lattice. (E) Left: Schematic of microtubule with closed and opened lattice. Right: Open microtubule fraction quantified as the total opened microtubule length divided by total observed microtubule length. Sample sizes from left to right on the graph: n=36, 21, 119, 61, where n is reported as number of TEM frames analyzed.

### Oxidative stress does not alter GTP-tubulin hydrolysis rate or GTP-tubulin lattice affinity

One way in which oxidative stress could increase rescue frequency would be by slowing the hydrolysis rate of stabilizing GTP-tubulin subunits within the microtubule lattice, perhaps as a consequence of cysteine oxidation. Here, because GTP-tubulin subunits are more stable in the microtubule lattice than their hydrolyzed GDP-tubulin counterparts, a slowing of the GTP-tubulin hydrolysis rate could increase the frequency of rescue events. To test this possibility, we incubated tubulin together with GTP at either 0°C or 37°C for 2.5 h, after which the reactions were stopped and inorganic phosphate release was measured using a CytoPhos Biochem Kit (Cytoskeleton, Inc.; BK054) (Portran et al., 2017). We found that, consistent with previous results, the hydrolysis rate of GTP was ∼67% higher at 37 °C, where microtubule polymerization occurs, relative to 0 °C, where microtubule polymerization is prevented (Portran et al., 2017). When 0.5 mM H_2_O_2_ was added to the reaction mixture, we found that the rate of GTP hydrolysis was unchanged relative to the controls, both at 0 °C and at 37 °C (Fig. 3B, n.s.=not significant, Mann-Whitney Test), indicating that oxidative stress does not alter the rate of GTP hydrolysis within polymerized microtubules.

Another potential explanation for increased rescue frequency could be that GTP-tubulin subunits are more stable within polymerized microtubules. One sensitive measure to gage the stability of GTP-tubulin subunits within a microtubule is to measure the nucleation time of new microtubules, which relies on GTP-tubulin subunits binding to each other, to start the growth of new microtubules. Thus, a fluorescence spectrophotometer was used to measure turbidity, a quantitative measure of bulk microtubule assembly (Fig. 3C, inset) (Portran et al., 2017). Turbidity was measured immediately after an ice-cold tubulin mixture was added to a pre-warmed 30 °C cuvette, and then over the following 30 min time interval, in the presence or absence of 0.5 mM H_2_O_2_ (Fig. 3C, inset). Microtubule nucleation time was quantified as the time delay prior to rapid tubulin polymerization. We found that the presence of H_2_O_2_ did not alter microtubule nucleation time (Fig. 3C). Thus, an oxidative stress-induced increase in microtubule rescue frequency is not due to a slowed hydrolysis rate of GTP-tubulin subunits within the microtubule lattice, nor due to an increase in stability of GTP-tubulin subunits within the microtubule itself.

### Electron microscopy reveals that oxidative stress causes structural damage to the GDP-Tubulin microtubule lattice

Because H_2_O_2_ exposure oxidizes cysteine residues within tubulin subunits, and tubulin cysteine oxidation has been previously implicated in weakened tubulin-tubulin interactions (Landino et al., 2002, 2011), we then tested whether oxidative stress could alter microtubule structure, perhaps by altering the stability of GDP-Tubulin subunits within the microtubule lattice. Thus, we used negative-stain transmission electron microscopy (TEM) to examine control and H_2_O_2_ treated microtubules at nanoscale resolution. Reconstituted GDP-tubulin microtubules were placed on a prewarmed 300-mesh carbon coated copper grid for 1 min, stained with uranyl acetate, and then the microtubules were imaged using TEM (FEI Technai Spirit BioTWIN). We observed that the H_2_O_2_-treated microtubules frequently had opened stretches, as well as smaller defects and holes, along the microtubule lattice (Fig. 3D). To quantify this observation, we characterized the fraction of open microtubule stretches relative to the total observed microtubule length (Fig. 3E, left). We found that there was a 4-fold (325%) increase in open microtubule sections in 0.5 mM H_2_O_2_ relative to untreated controls (Fig. 3E, right; fit to exponential equation as shown, p=0.999, Χ^2^ test statistic=0.0028). Thus, oxidative stress acts to disrupt GDP-tubulin stability within the microtubule lattice, leading to nanoscale structural defects within the microtubule itself (Fig. 3E).

### Oxidative stress-induced structural defects are repaired via the incorporation of GTP-tubulin subunits into the GDP-tubulin lattice

Paradoxically, we found that oxidative stress increases microtubule rescue frequency, leading to longer microtubules, but also disrupts GDP-tubulin interactions within the microtubule, leading to opened sections and defects within the microtubule itself. How could openings and holes within the microtubule lead to increased rescue frequency and thus net elongation of dynamic microtubules? Interestingly, recent publications have shown that, in the presence of free tubulin, open defects within a microtubule can be “repaired” via the incorporation of GTP-tubulin subunits into the microtubule itself, both in cells and in purified systems (Aumeier et al., 2016; Dimitrov et al., 2008; de Forges et al., 2016; Schaedel et al., 2015, 2019). Further, it has been shown that the incorporation of GTP-tubulin at damaged areas within the GDP-tubulin lattice acts to stabilize the microtubule in these locations, leading to an increased frequency of rescue events at the “repaired” site (Fig. 2A) (Aumeier et al., 2016; Dimitrov et al., 2008; de Forges et al., 2016; Tropini et al., 2012).

Thus, we tested whether oxidative stress could promote incorporation of GTP-tubulin into the GDP-tubulin microtubule lattice, due to openings within the microtubule itself (Fig. 4B). We used our previously published “repair assay” to visualize the incorporation of soluble Alexa488-labeled green GTP-tubulin into control and H_2_O_2_ treated rhodamine-labeled (red), GDP-tubulin microtubules (Fig. 4B) (Reid et al., 2017). Here, GDP-tubulin microtubules were treated for 1h in buffer alone, or in 0.5 mM H_2_O_2_, and then the microtubules were mixed with a solution containing Alexa488-labeled, green GTP-tubulin (see materials and methods). After a 1h incubation, microtubules were imaged using TIRF microscopy. The incorporation of green GTP-tubulin within the red microtubules was observed as green puncta along the length of the red microtubules (Fig. 4C, left, yellow arrows). To quantify the incorporation of GTP-tubulin within the microtubule itself, we measured the fractional coverage area of green tubulin along the red microtubule lattice (Fig. 4C, left, yellow arrows), excluding green tubulin at the microtubule ends, which represents normal microtubule elongation (Fig. 4C, left, white arrows). We found that there was a 94% increase in incorporation of GTP-tubulin along the length of the microtubule for H_2_O_2_ treated microtubules, relative to untreated controls (2-fold, p<<0.0001, Student’s t-test). This rate of incorporation was less than the 4-fold increase in open microtubule sections observed via electron microscopy (Fig. 3E), suggesting that GTP-tubulin does not completely incorporate into all of the damaged areas within the microtubule lattice.

**Figure 4:**
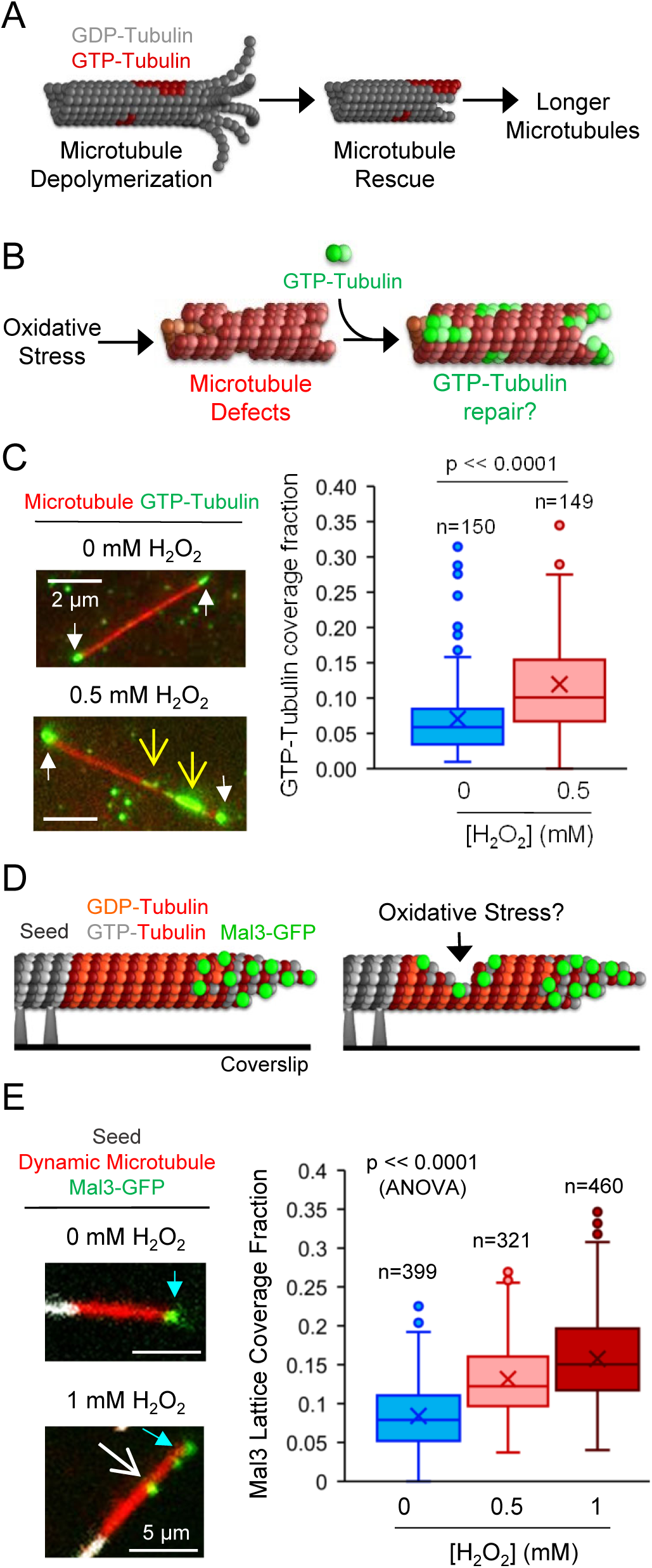
Oxidative stress leads to damage in the GDP-tubulin microtubule lattice that is repaired via the incorporation of GTP-tubulin subunits. (A) Incorporation of GTP-tubulin into the GDP-tubulin lattice (left) leads to stabilized areas within the GDP-tubulin lattice (middle), increasing the likelihood of rescue events, and thus longer microtubules (right) (Aumeier et al. 2016; Dimitrov et al. 2008; de Forges et al. 2016; Gardner 2016; Tropini et al. 2012). (B) Schematic of microtubule repair assay to measure the incorporation of GTP-tubulin (green) into damaged areas on stabilized GDP-tubulin microtubules (red). (C) Left: TIRF microscopy images of GFP-Tubulin (green) and GDP-tubulin microtubules (red). White arrows: microtubule growth via addition of GTP-tubulin to microtubule ends (excluded from analysis). Yellow arrows: incorporation of GTP-tubulin (green) into damaged areas on the GDP-tubulin lattice (red). Right: Quantification of the coverage area of GTP-tubulin (green) lattice incorporation divided by the total GDP-tubulin microtubule lattice area (red). Green tubulin growth from microtubule ends is excluded from the analysis. Crosses: Mean; Bar: Median; Box: first-third quartiles; sample sizes represent number of images, many microtubules per image. (D) Schematic of Mal3-GFP Binding Assay. Left: In a dynamic microtubule assay, Mal3-GDP typically binds to growing microtubule ends. Right: Mal3-GFP may bind to damaged areas on the lattice, into which GTP-tubulin has incorporated. (E) Left: Representative TIRF images of Mal3-GFP binding assay, in which Mal3-GFP (green) binds the dynamic microtubule extension (red) polymerized from stabilized seeds (white). Mal3-GFP binds to growing microtubule tips, as expected (cyan arrows), but can also be observed within the microtubule lattice (white arrow), especially in the presence of H_2_O_2_ (bottom). Right: Quantification of Mal3-GFP coverage fraction on microtubules. Crosses: Mean; Bar: Median; Box: first-third quartiles; sample sizes represent number of images, many microtubules per image.

### Oxidative stress-induced structural defects are repaired via the incorporation of GTP-tubulin subunits into the GDP-tubulin lattice during the short lifetime of dynamic microtubules

Both the electron microscopy experiments and the GTP-tubulin repair experiments described above were performed by treating stabilized GDP-tubulin microtubules with H_2_O_2_ for 60 minutes or more. To verify that oxidative stress mediated damage and GTP-tubulin repair could occur over the much shorter lifetime of dynamic microtubules in cells (lifetime 3-5 minutes in our cell-free assay, Fig. 1D), we used dynamic microtubules and a microtubule tip-tracking protein, Mal3-GFP. Mal3 and its human homologue EB1 have been shown to bind with high affinity to GTP-tubulin, and with lower affinity to GDP-tubulin (Maurer et al., 2011; Zanic et al., 2009). Further, EB1 has been shown to bind rapidly to microtubule structural defects, increasing the efficiency at which EB1 (or Mal3) may detect and stably bind to GTP-tubulin that is associated with openings and holes within the microtubule itself (Reid et al., 2019). Thus, to determine whether oxidative stress-mediated damage and GTP-tubulin repair occurs in short-lived dynamic microtubules, we examined the binding of Mal3-GFP to dynamic microtubules in the presence of increasing H_2_O_2_ concentrations.

To perform the dynamic microtubule assay in the presence of Mal3-GFP, Alexa647-labelled (far-red) dynamic microtubules (Fig. 4D, red) were grown from rhodamine-labeled seeds (Fig. 4D, grey). Using this system, Mal3-GFP is expected to be visible in the green channel at microtubule tips (Fig. 4D, left, green; Fig. 4E, left, cyan arrows). In the presence of microtubule damage and repair, we predicted that Mal3-GFP would also be observed along the length of the microtubule, in addition to the growing microtubule tip (Fig. 4D, right; Fig. 4E, left, white arrow).

As expected, Mal3-GFP was observed at growing microtubule tips (Fig. 2D, left-top), and less frequently along the microtubule length (Fig. 2D, left-bottom). To quantify the relative amounts of Mal3-GFP binding in the presence and absence of oxidative stress, we measured the area of green Mal3-GFP occupancy along the length of each microtubule, divided by the total area of the red microtubules, for each image (Reid et al., 2017, 2019). We found that the coverage of Mal3-GFP along the length of dynamic microtubules was significantly and substantially increased with increasing H_2_O_2_ concentration (Fig. 2D, right, ANOVA for comparison between groups, p<<0.0001; sample sizes indicate number of images analyzed, each of which contain many microtubules). Further, there was a 90% increase in binding of Mal3-GFP to dynamic microtubules in 0.5 mM H_2_O_2_ relative to the controls (p<<0.0001, t-test), which was nearly identical to the 94% increase in GTP-tubulin repair in the stable microtubules, suggesting that oxidative stress acts on a rapid time scale (< 5 minutes) to damage microtubules, allowing for swift GTP-tubulin repair at these sites.

### Oxidative stress leads to acetylation of cardiac myocyte microtubules

It has been previously shown that the presence of openings and holes along the microtubule length also increases the likelihood of post-translational microtubule acetylation, by permitting access of the acetylation enzyme αTAT1 to its acetylation site inside the hollow lumen of the microtubule (Coombes et al., 2016; Janke and Montagnac, 2017; Ly et al., 2016). Thus, we asked whether the lengthened, longitudinally aligned, and likely damaged microtubules that were generated under conditions of oxidative stress in cardiac myocytes were also acetylated.

To test this idea, rat ventricular cardiac myocytes were treated with 0, 100, or 500 μM H_2_O_2_ for 1h at 37 °C. The cover slip-adherent cardiac myocytes were then washed with PBS, fixed in 1% PFA, stained with anti-acetylated-tubulin antibody (6-11B), and imaged with confocal microscopy (Fig. 5A).

**Figure 5:**
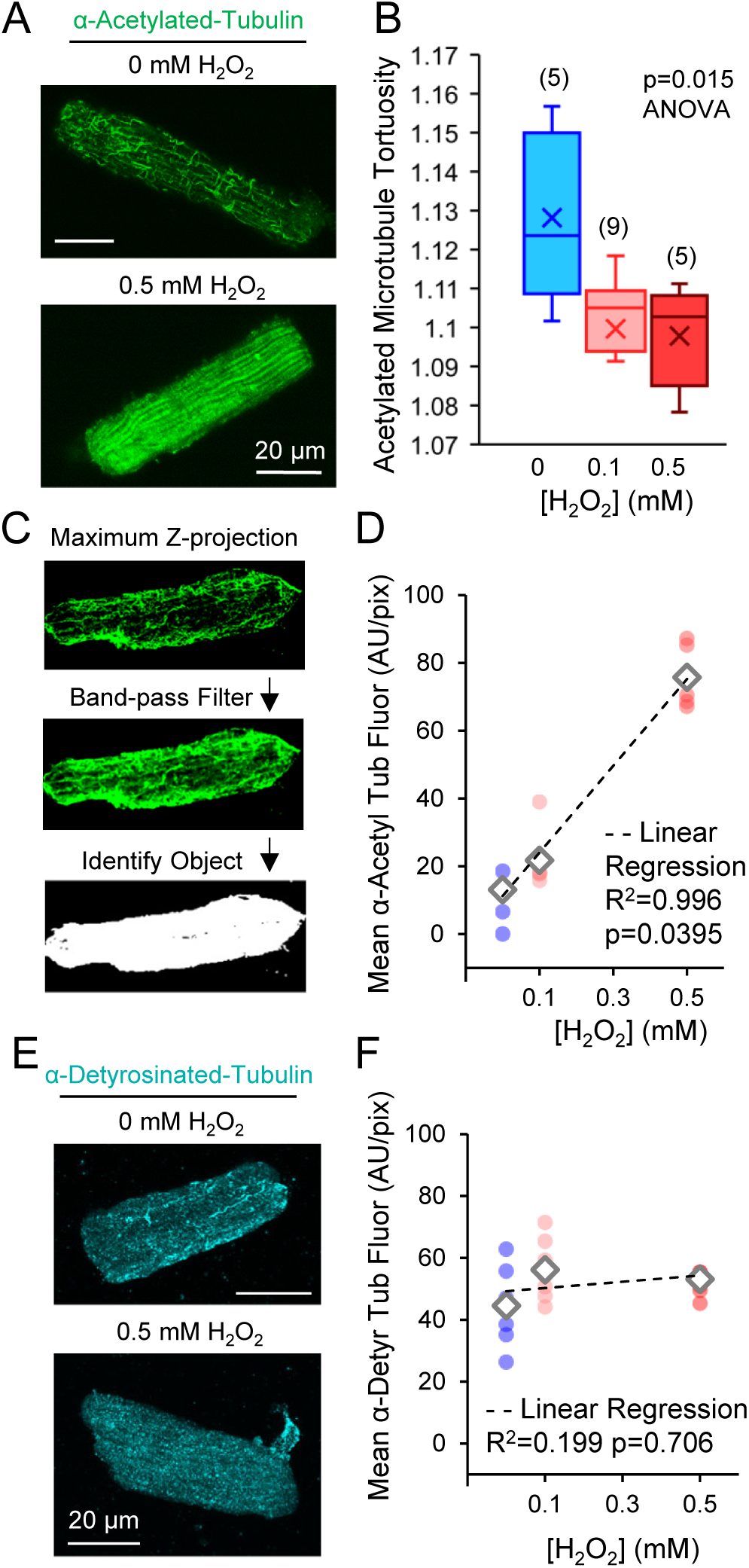
In cardiac myocytes exposed to oxidative stress, longitudinally aligned microtubules are acetylated. (A) Rat ventricular cardiac myocytes fixed and stained for anti-acetylated tubulin (green). (B) Quantification of mean acetylated microtubule tortuosity. Crosses: Mean; Bar: Median; Box: first-third quartiles; sample sizes represent number of cells analyzed, many microtubules per cell. (C) Method for using automated object detection to identify cardiac myocytes and then measure average anti-acetylated tubulin fluorescence intensity per pixel inside of each cell. A maximum Z-projection image is used for analysis, to ensure that all fluorescence within the cell is captured in the analysis. (D) Quantification of mean anti-acetylated tubulin fluorescence intensity per pixel in cardiac myocytes. Each circle marker represents one image, n=5 images per group; open diamonds represent average values for each group, dotted line shows best-fit linear regression for average values. (E) Rat ventricular cardiac myocytes fixed and stained for anti-detyrosinated tubulin (cyan). (F) Quantification of mean anti-detyrosinated tubulin fluorescence intensity per pixel in cardiac myocytes. A maximum Z-projection image is used for analysis, to ensure that all fluorescence within the cell is captured in the analysis. Each circle marker represents one image, n=7, 10, 8 images per group from left to right on the graph; open diamonds represent average values for each group, dotted line shows best-fit linear regression for average values.

First, to determine whether acetylation was prevalent on the lengthened, longitudinally aligned microtubules generated via oxidative stress, we measured the mean tortuosity of acetylated microtubules in cardiac myocytes exposed to oxidative stress. Similar to results for α-tubulin staining, we found that the mean acetylated microtubule tortuosity was significantly decreased with exposure to oxidative stress (Fig. 5B, p=0.015 ANOVA, difference between groups, sample sizes=number of cells analyzed), suggesting that the lengthened, longitudinal microtubules that are prevalent in cardiac myocytes under conditions of oxidative stress are also acetylated.

Second, we tested whether the overall microtubule acetylation level was increased in cardiac myocytes exposed to oxidative stress. Thus, we used automated object detection code in MATLAB to identify cardiac myocytes within our anti-acetylated-tubulin antibody images (Fig. 5C), and then used the code to measure the average anti-acetylated tubulin fluorescence intensity per pixel within each cardiac myocyte, using a maximum Z-projection images to capture all of the fluorescence throughout the depth of the entire cardiac myocyte (Fig. 5C, top). We found that the mean anti-acetylated tubulin fluorescence intensity increased 5.8-fold (485%) in cardiac myocytes treated with 500 μM H_2_O_2_ relative to controls (Fig. 5D, linear regression for slope, p=0.0395).

To distinguish whether the increased overall acetylation level was an indirect consequence of altered microtubule mass, or whether microtubule acetylation was directly facilitated by oxidative stress, we measured the overall level of an alternative microtubule post-translational modification, detyrosination (Zhou et al., 2019). Detyrosination occurs on the outside of the microtubule, and so we predicted that the rate of microtubule detyrosination would not be altered by the presence of openings and holes along the microtubule. To test this idea, rat ventricular cardiac myocytes were treated with 0, 100, or 500 μM H_2_O_2_ for 1h at 37 °C. The cover slip-adherent cardiac myocytes were then washed with PBS, fixed in 1% PFA, stained with anti-detyrosinated-tubulin antibody, and imaged with confocal microscopy. In contrast to the anti-acetylated tubulin antibody, we found no significant increase in detyrosinated tubulin signal with H_2_O_2_ exposure (Fig. 5F, 5G, linear regression for slope, p=0.706). This result suggests that acetylation of microtubules was a direct outcome of cardiac myocyte exposure to oxidative stress, and not an indirect effect of altered microtubule mass.

## Discussion

In this work, we found that reactive oxygen species, which are generated under conditions of oxidative stress, oxidize microtubules (Fig. 3A). To our knowledge, this is the first demonstration of direct oxidation of polymerized microtubules, in the absence of glycolytic enzymes or other microtubule associated proteins, via oxidative stress. Our results indicate that exposure to reactive oxygen species leads to the irreversible cysteine oxidation of tubulin subunits within a microtubule. Tubulin subunit cysteine oxidation has been implicated in weakened tubulin-tubulin interactions, and thus likely increases the susceptibility of microtubules to damage (Clark et al., 2014; Guo et al., 2006; Landino et al., 2002, 2011; Luduena and Roach, 1991; Mellon and Rebhun, 1976) Consistent with this implication, we found that oxidized microtubule were characterized by regions of damage along the length of the microtubule, specifically, by holes and open sections within the GDP-tubulin microtubule lattice (Fig 3). This oxidation and damage occurs on a rapid time scale (< 5 minutes) (Fig. 4), thus leading to significant alterations in the behavior of dynamic microtubules (Fig. 2).

Paradoxically, oxidative-stress mediated damage along the length of the microtubule results in the lengthening of dynamic microtubules, via an increase in the frequency of rescue events (Fig. 2). Rescue frequency is increased as a result of GTP-tubulin incorporation at damaged microtubule lattice sites (Aumeier et al., 2016; Dimitrov et al., 2008; Dye et al., 1992; de Forges et al., 2016; Schaedel et al., 2015, 2019). As previously described, sites of GTP-tubulin incorporation along the length of the microtubule can slow down and ultimately stop microtubule depolymerization, allowing the microtubule to begin growing again (Aumeier et al., 2016; Dimitrov et al., 2008; de Forges et al., 2016; Tropini et al., 2012). The increased rescue frequency of dynamic microtubules upon exposure to oxidative stress led to an exponential increase in predicted dynamic microtubule lengths. These results are consistent with the recent finding that microtubule severing enzymes are able to increase microtubule length and amplify the cytoskeleton by causing nanoscale damage along the microtubule length, leading to GTP-tubulin islands that stabilize the microtubules against depolymerization(Vemu et al., 2018).

Importantly, in cardiac myocytes, the lengthening of dynamic microtubules increases the likelihood of their direct interaction and binding to regularly spaced Z-disks (Fig. 6, top, red: dynamic microtubules, blue: Z-disks) (Robison et al. 2016; Robison and Prosser 2017). It has been previously shown that the binding of microtubules to Z-disks inhibits microtubule depolymerization (Drum et al. 2016). Thus, the structure of the cardiac myocyte itself exacerbates the effects of oxidative stress: lengthened dynamic microtubules are likely to be stabilized against depolymerization when they grow long enough to interact with the regularly spaced Z-disks (Fig. 6, bottom). This model of dynamic microtubule lengthening, combined with binding of lengthened microtubules to the Z-disk, predicts that oxidative stress generates a longitudinally aligned and static microtubule network in cardiac myocytes, likely increasing cellular stiffness and disrupting contractility (Fig. 6, bottom). Consistent with this model, Drum et al. observed a reduction in dynamic microtubule tips, as evidenced by the absence of EB3-GFP microtubule growth and shortening events, after 30 min of H_2_O_2_ exposure (Drum et al., 2016). This loss of dynamic microtubule tips under conditions of oxidative stress is suggestive of a shift in the balance between static and dynamic microtubules, as the lengthened dynamic microtubules are stabilized over time via interactions with the Z-disk (Fig. 6, bottom).

**Figure 6:**
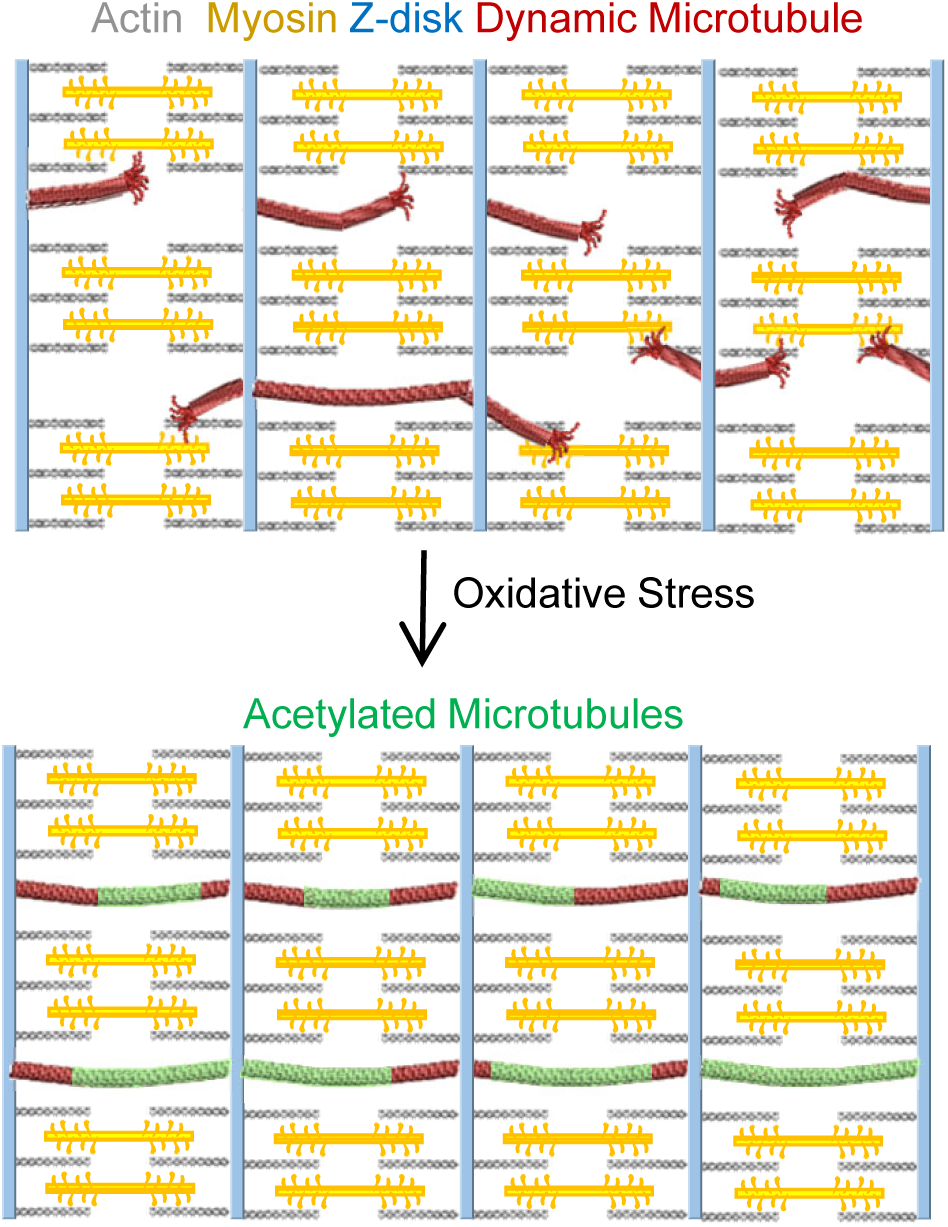
Model for oxidative stress mediated remodeling of the cardiac myocyte microtubule cytoskeleton. Model for longitudinal realignment of cardiac myocyte microtubules with oxidative stress: the switch from a sparse, complex microtubule network (top) into a longitudinally oriented network (bottom) suggests that lengthened dynamic microtubules (red) may be aligned and stabilized by interaction and binding of the growing microtubule plus-ends with intermediate filaments at the Z-disks (blue) (Robison et al., 2016). In addition, oxidative-stress mediated damage to microtubules facilitates extensive acetylation of the cardiac myocyte microtubules (bottom, green).

The effect of oxidative stress in damaging microtubules has a further consequence, as the damaged microtubule lattice fosters a change in the post-translational acetylation state of microtubules. Specifically, defects, gaps, and holes along the microtubule lattice have been shown to allow improved access of the acetylation enzyme αTAT1 to its acetylation site inside the hollow lumen of the microtubule, thus increasing the likelihood of microtubule acetylation (Coombes et al., 2016; Janke and Montagnac, 2017; Ly et al., 2016). Indeed, we observed a dramatic increase in microtubule acetylation in oxidative-stress treated cardiac myocytes (Fig. 5; Fig. 6, bottom). Importantly, acetylated microtubules are resistant to breakage, which increases their longevity (Eshun-Wilson et al., 2019; Portran et al., 2017; Xu et al., 2017). This increased microtubule longevity likely further reinforces the oxidative stress-mediated transformation from a versatile dynamic microtubule network into a longitudinally aligned and static microtubule network in diseased cardiac myocytes.

Previous reports have been contradictory regarding the effect of oxidative stress and ischemia on microtubule density inside of cells (Drum et al., 2016; Hinshaw et al., 1993; Kratzer et al., 2012; Valen et al., 1999; Wei-Guo and Qi-Ping, 2014). We did not find a significant effect of oxidative stress on the rate of microtubule nucleation, and so we posit that the most important effect of oxidative stress in cardiac myocytes is on the lengthening, longitudinal reorientation, and acetylation of microtubules, rather than on a direct increase in microtubule number (Fig. 6). Specifically, by increasing both the length and the lifetime of microtubules, oxidative stress could act to disrupt cardiac myocyte contractility, regardless of an increase in total numbers or density of microtubules.

Previous work has implicated the detyrosination of α-tubulin in cardiac myocyte contractile dysfunction and patient heart failure (Chen et al., 2018; Kerr et al., 2015; Robison et al., 2016). In the present study, oxidative stress itself did not lead to an increase in cardiac myocyte microtubule detyrosination (Fig. 5F,G). However, microtubule detyrosination has been reported to increase the affinity of microtubules to Z-disk intermediate filaments such as Desmin (Robison et al., 2016), which could potentially have an additive effect in increasing cardiac myocyte stiffness when combined with the direct effects of oxidative stress in patients with heart failure.

Our single microtubule studies in cell-free experiments permit high resolution measurements of the direct effect of H_2_O_2_ on dynamic microtubules. However, we cannot exclude the possibility that microtubule associated proteins and other factors inside of cells could have a confounding effect on microtubule dynamics, over and above the direct effect of H_2_O_2_ as was assessed in our cell-free assay. For example, previous work suggests that oxidative stress causes displacement of the microtubule-associated protein EB1 from growing microtubule ends, which could contribute to differences in microtubule dynamics in cells as compared to our cell-free assay (Le Grand et al., 2014; Smyth et al., 2010).

Regardless, we find that oxidative stress acts inside of cardiac myocytes to facilitate a dramatic, pathogenic shift from a dynamic, versatile microtubule network into a highly acetylated, longitudinally aligned, and static microtubule network (Fig. 6), which likely contributes to increased cellular stiffness and contractile dysfunction in failing heart cells. This deleterious effect of oxidative stress in contractile cells may have important broader health impacts, such as in muscle dystrophies, ischemic heart disease, systolic and diastolic heart failure, and a whole host of cardiomyopathies.

## Materials and Methods

### Ventricular Cardiac myocyte Isolation and Primary Culture

Adult rat ventricular myocytes were isolated as previously described (Thompson et al., 2016, 2019). Briefly, adult female rats were anaesthetized (isoflurane) and injected with heparin (15,000 U/kg) and Fatal Plus (150 mg/kg). Enzymatic digestion of hearts was achieved via the Langendorff procedure. Ventricles were triturated and cells were plated on laminin-coated glass coverslips and cultured overnight at 37°C in M199 media (Sigma) supplemented with 25mM HEPES, 26.2mM sodium bicarbonate, 0.02% BSA, 50U/mL penicillin-streptomycin, 5ug/mL insulin, 5ug/mL transferrin, and 5ng/mL selenite and pH-adjusted to 7.4.

### H_2_O_2_ exposure, immunostaining, and imaging of cardiac myocytes

Coverslip-adherent cardiac myocytes were treated with pre-warmed H_2_O_2_ (0-1mM) in M199 media for 1 h at 37°C, washed in pre-warmed PBS, and fixed by incubating in 1% paraformaldehyde in pre-warmed PBS for 1h. Coverslips were blocked in blocking buffer containing 3% BSA and 0.1% Triton X-100 in PBS for 30min at room temperature. For immunostaining of α-tubulin, coverslips were probed with mouse anti-α-tubulin antibody (DM1A, VWR; #PI62204; 1:1000 in blocking buffer) for 1h at room temperature, washed twice in blocking buffer for 10min each, and treated with FITC anti-mouse IgG antibody (F0257, Sigma; 1:1000 in blocking buffer) overnight at 4°C. For immunostaining of acetyl-tubulin, coverslips were probed as above, but with mouse anti-acetyl-tubulin antibody (T7451, Sigma; 1:2000 in blocking buffer) for 1h at room temperature, washed twice in blocking buffer for 10min each, and treated with FITC goat anti-mouse antibody (F0257, Sigma; 1:2000 in blocking buffer) overnight at 4°C. For immunostaining of detyrosinated tubulin, coverslips were probed with rabbit anti-detyrosinated tubulin antibody (AB3201, Sigma; 1:1000 in blocking buffer) for 1h at room temperature, washed twice in blocking buffer for 10min each, and treated with donkey Alexa Fluor 488-conjugated anti-rabbit IgG secondary antibody (A21206, Life Technologies; 1:1000 in blocking buffer) overnight at 4°C. Coverslips were again washed twice in blocking buffer for 10min and mounted on slides with ProLong Diamond Antifade (Thermo Fisher #P36965) for imaging. Immunostained cardiac myocytes were imaged with a laser scanning confocal microscopy (Nikon Ti2, 488nm laser line) fitted with a 100x oil objective (Nikon N2 Apochromat TIRF 100X Oil, 1.49 NA), which allowed for a 0.2µm pixel size.

### Microtubule Tortuosity Measurement in Cardiac myocytes

Quantification of microtubule tortuosity was performed using custom-made automated MATLAB script. Immunostained cardiac myocyte images were filtered for noise and background with Gaussian band-pass filtering. Microtubules were identified using Canny edge detection to create a skeleton map. The skeleton map was segmented at branch points to create microtubule segments, which were filtered to exclude segments shorter than 7 pixels. Quasi-Euclidean distance (path length) and Pythagorean distance (endpoint displacement) were measured for each microtubule segment. Tortuosity was calculated by dividing path length by endpoint displacement for each microtubule segment, and mean tortuosity value for each image was calculated across all microtubule segments.

### Tubulin Purification and Labeling

Tubulin was purified from pig brain extract through repeated cycles of polymerization-depolymerization and labeled with rhodamine, Alexa-488, or Alexa-647 as previously described (Castoldi and Popov, 2003; Gell et al., 2010).

### Construction and preparation of flow chambers for TIRF microscopy imaging

Flow chambers were assembled for TIRF microscopy as described (Gell et al., 2010) with the following adaptation: To create a ‘lane’ for the unidirectional flow of samples, two narrow strips of Parafilm were arranged parallel to each other in between two hydrophobic silanized coverslips. Chambers were subjected to heat in order to melt the Parafilm strips and create a seal between the coverslips. Before use, the chamber was treated with rabbit anti-rhodamine antibody diluted 1:50 (A6397, Thermo Fisher) in Brb80 for 30 min followed by blocking with pluronic F127 for at least 20 min.

### Construction of stabilized microtubule seeds

Stable GMPCPP microtubule seeds were prepared from a 40.5 µL mixture of 3.9 µM tubulin (25% rhodamine-labeled, 75% unlabeled), 1mM GMPCPP, and 1.2 mM MgCl_2_ in Brb80 and incubated first on ice for 5 min, followed by 2 h at 37°C (Reid et al., 2017). Following incubation, GMPCPP microtubules were diluted in 400 µL warm Brb80, spun via air-driven centrifuge (Airfuge, Beckman Coulter, 20psi, 5min), and resuspended in 400 µL of warm 10 µM Taxol in Brb80. These GMPCPP microtubules were stored at 37°C and used for microtubule dynamics experiments up to 5 days after preparation.

### Dynamic Microtubule Assay and TIRF Microscopy

Rhodamine-labeled GMPCPP microtubule ‘seeds’ were adhered to an anti-rhodamine antibody-coated flow chamber (Gell et al., 2010) and washed with 80 µL pre-warmed Imaging Buffer (20µg/mL glucose oxidase, 10µg/mL catalase, 20mM D-Glucose, 10mM DTT, 80µg/mL casein, and 1% tween-20) to minimize photobleaching. A reaction mixture containing 10.3µM tubulin (15.8% Alexa488-labeled, 84.2% unlabeled), 55mM KCl, 1 mM GTP, 0.15% methyl cellulose and Imaging Buffer was prepared. For oxidative stress experiments, H_2_O_2_ (0-1000 µM) was included in the reaction mixture. The reaction mixture was centrifuged for 5 min at 4°C to remove protein aggregates and the supernatant was introduced into the imaging chamber.

Movies of dynamic microtubules were acquired at 28°C for 1h at 0.2fps using 488nm and 561nm laser lines with a TIRF microscope (Nikon Eclipse Ti TIRF) fitted with an 100x oil objective (Nikon CFI Apochromat TIRF 100XC Oil, 1.49 NA) and CCD camera (Andor, iXon3). This TIRF microscopy imaging system allowed for a 160nm pixel size. With this experimental protocol, multiple parameters of length dynamics (elongation rate, shortening rate, rescue frequency, and catastrophe frequency) were quantified using ImageJ (Coombes et al., 2013; Gardner et al., 2011b, 2011a; Reid et al., 2016).

### Dynamic Microtubule Assay Image Analysis

Kymographs for each dynamic microtubule extension were generated from movies using ImageJ. For each microtubule, the growth rates, shortening rates, catastrophe frequency, and rescue ratio was quantified. For the growth rate analysis, only microtubules in which the entire lifetime of microtubule elongation was observed were analyzed. Growth rate was calculated as the microtubule length at catastrophe divided by the duration of microtubule growth, from growth initiation from the seed until catastrophe. For the catastrophe frequency analysis, the catastrophe frequency was calculated as the inverse of the duration of microtubule growth. For the shortening rate analysis, only complete shortening events, defined as the period following catastrophe until either complete depolymerization or rescue, were analyzed. Shortening rate was calculated as the length a microtubule shortened after catastrophe divided by the shortening time. For the rescue frequency analysis, because microtubule rescue only occurs after a catastrophe event, the probability of rescue is first dependent on a catastrophe event. Therefore, the rescue frequency was calculated as number of rescue events in a movie, divided by the number of catastrophe events, and then normalized to the mean microtubule length at catastrophe

### Microtubule Turbidity Nucleation Assay

Bulk tubulin polymerization was assessed by measuring sample turbidity, as the absorbance at 350 nm (Nanophotometer NP80, Implen), as previously described (Portran et al., 2017). Briefly, the spectrophotometer was pre-warmed in a 30°C room for 2h before use, and blanked on dH_2_O. A mixture of 42µM tubulin, 1mM GTP, 5mM DTT, and 5% glycerol was prepared in Brb80 on ice. For H_2_O_2_ experiments, this mixture additionally contained 0.5mM H_2_O_2_. The tubulin mixture was added to a pre-warmed cuvette and absorbance measurements were acquired every 30sec for 30min.

### GTP Hydrolysis assay

A mixture of 10 µM tubulin, 0.5mM GTP, 0.5mM DTT, and 5% glycerol in Brb80 was prepared on ice. For H_2_O_2_ experiments, this mixture also included 0.5 mM H_2_O_2_. Reactions were incubated either at 0°C or 37°C for 2.5h, after which reactions were stopped by the addition of 5% TCA on ice for 2min. Samples were centrifuged for 10min at 4°C to remove protein aggregates. 15µL of the supernatant was added to 35µL Cytophos reagent (BK054, Cytoskeleton), vortexed gently, and incubated for 10min at 22°C. To measure the amount of inorganic phosphate, the absorbance at 650nm (Nanophotometer NP80, Implen) was recorded three times per sample. A phosphate standard curve was determined using the Cytophos phosphate standard (BK054, Cytoskeleton), diluted 0-100µM.

### Measurement of Tubulin Cysteine Oxidation

A mixture of 36.4 µM tubulin (12% Alexa488-labeled, 21% biotinylated), 1mM GTP, and 315µM DCP-Rho1 was prepared on ice and incubated for 2h at 37°C in the presence or absence of 1mM H_2_O_2_. The mixture was diluted 100-fold with warm 10µM Taxol in Brb80 and stored overnight at 37°C. 210µL of the microtubule sample were spun via air-driven centrifuge (20psi, 5min), and re-suspended in 150µL of warm 10µM Taxol in Brb80. An imaging chamber was assembled as described above with the following variation: Instead of anti-rhodamine antibody, imaging chambers were treated with a mixture of 5µL rabbit anti-biotin antibody (D5A7, company), 20µL streptavidin (1mg/mL), and 15µL Brb80 for 30min, before blocking with pluronic F127. 80µL warm 10µM Taxol in Brb80 was first flushed through the imaging chamber followed by 40µL of the Taxol-stabilized microtubules. Non-adherent microtubules were removed by flushing with warm 10µM Taxol in Brb80. Imaging Buffer was introduced and microtubules were imaged via TIRF microscopy, as described above.

### Transmission Electron Microscopy (TEM)

A mixture of 37µM tubulin, 1mM GTP, 4.3mM MgCl_2_, and 4% DMSO was prepared and placed on ice for 5min, followed by a 30min incubation at 37°C. Reconstituted microtubules were diluted 20x in warm 20µM Taxol in Brb80. Taxol-stabilized microtubules were diluted 2x in a pre-warmed solution containing H_2_O_2_ or Brb80 alone, and stored overnight at room temperature. A drop of reconstituted microtubules was placed on a pre-warmed 300-mesh carbon coated copper grid for 1min, followed by 1 drop of 1% uranyl acetate for 1min. Filter paper was then used to wick away the excess stain from the grid. The grid was left to dry for 10min and stored. Microtubule specimens were imaged using TEM (FEI Technai Spirit BioTWIN).

### Microtubule Repair Assay

GDP microtubules were prepared from a mixture of 34.8µM tubulin (25% rhodamine-labeled, 75% unlabeled), 4.3mM MgCl2, 1mM GTP, and 4.3% DMSO in Brb80. The mixture was incubated for 1h at 37°C, diluted 10x in warm 10µM Taxol in Brb80, and stored overnight at 37C. 250µL of the Taxol-stabilized microtubules were then spun via air-driven ultracentrifuge (20psi, 5min), resuspended in 50µL of warm 10µM Taxol in Brb80 with or without H_2_O_2_, and incubated for 30min at 37C. To visualize the incorporation of soluble tubulin at ‘defects,’ or gaps, within the microtubule lattice, a reporter tubulin was used (Reid et al., 2017). Briefly, 20µL of the resulting microtubule mixture was added to 20µL of tubulin reporter solution containing 1.5µM tubulin (50% Alexa488-labelled), 1mM MgCl_2_, 250µM GTP, and 10µM Taxol in Brb80 and incubated for 30min at 37°C. Microtubules were then adhered to an imaging chamber and imaged via TIRF microscopy as described above.

### Microtubule Repair Assay Image Analysis

To quantify incorporation of green GTP-tubulin, the fractional coverage area of green tubulin along the red microtubule was determined using a previously described custom built MATLAB analysis tool (Reid et al., 2017). First, the red microtubule channel was processed to identify GDP microtubules. The green reporter tubulin channel was then filtered to reduce high-frequency noise and smoothing was performed to correct for heterogeneity within TIRF illumination of the optical field. A green channel threshold was manually selected just above background and maintained for consistent analysis across all experiments. For each TIRF image, the fractional area of green reporter tubulin incorporation within the red microtubule was recorded as the ‘Repair Fraction’, excluding green tubulin at the microtubule ends, which represents normal microtubule elongation.

### Mal3-GFP Purification

The pETMM11-HIS6x-Mal3-GFP plasmid with a TEV cut site after the His6x tag was a kind gift from Dr. Thomas Surrey. The plasmid was transformed into Rosetta (DE3) pLysS E. coli and grown in 800mL of LB+kan+cam at 37°C to an OD of approximately 0.4. To induce protein expression, IPTG was added to 0.2mM and the culture was mixed at 14°C for 16hr. Cells were centrifuged (30min., 4°C, 4400xg) and resuspended in 25mL lysis buffer (50mM Tris pH7.5, 200mM NaCl, 5% glycerol, 20mM imidazole, 5mM β-mercaptoethanol, 0.2% triton X-100), protease inhibitors (1mM PMSF, 10µM Pepstatin A, 10µM E-64, 0.3µM aprotinin), and DNAse I (1U/mL). The cell suspension was sonicated on ice (90% power, 50% duty, 6×1min). Cell lysates were centrifuged (1h, 4°C, 14000 xg) and the soluble fraction was passed through 1mL of Talon Metal Affinity Resin (Clontech #635509). The resin was washed for four times with 4mL Wash Buffer (50mM Tris pH7.5, 500mM NaCl, 5% glycerol, 20mM imidazole, 5mM β-mercaptoethanol, 0.1mM PMSF, 1µM Pepstatin A, 1µM E-64, 30nM aprotinin) for 5min each. Protein was eluted from the resin by mixing with 1mL of Elution Buffer (50mM Tris pH 7.5, 200mM NaCl, 250mM imidazole, 0.1mM PMSF, 1µM Pepstatin A, 1µM E-64, 30nM aprotinin) for 15min followed by slow centrifugation through a fritted column to retrieve eluate. To cleave the HIS6x tag, 10 units of GST-tagged TEV enzyme (TurboTEV, #T0102M, Accelagen) and 14mM β-mercaptoethanol were added and the eluate was dialysed into Brb80 overnight at 4°C. To remove the TEV enzyme, the dialysate was mixed with 100ul of glutathione-sepharose (GE Healthcare #17-0756-01) for 30 min. at 4°C and spun (1min, 2000xg). The Mal3-GFP protein was quantified by band intensity on a coomassie-stained SDS PAGE protein gel.

### Mal3 Microtubule Binding Assay

The microtubule dynamics assay was completed as described above with the reaction mixture containing 10µM tubulin (12.7% Alexa647-labeled, 87.3% unlabeled), 1mM GTP, 90mM KCl, 39.4nM Mal3-GFP, imaging buffer, and H_2_O_2_ (0 or 0.5mM). Images of dynamic microtubules were acquired at 28°C using 488nm and 561nm lasers with a TIRF microscope (Nikon Eclipse Ti TIRF) fitted with an 100x oil objective (Nikon CFI Apochromat TIRF 100XC Oil, 1.49 NA), CCD camera (Andor, iXon3) and, 2.5x projection lens. This TIRF microscopy imaging system allowed for a 64nm pixel size.

### Mal3 Microtubule Binding Assay Image Analysis

To compare relative binding of Mal3-GFP to dynamic microtubules in the presence and absence of H_2_O_2_, the Mal3 Coverage Fraction was calculated as the total area of green (Mal3-GFP) occupancy divided by the total area of the red microtubules for each image. A semi-automated MATLAB analysis code was used as previously described (Reid et al., 2017).

### Amplex Red assay to measure residual H_2_O_2_ in Imaging Buffer

In order to assess the concentration of H_2_O_2_ in the presence of the reactive oxygen species scavengers within imaging buffer, we used the previously established Amplex Red assay (Schlieve et al., 2006). Briefly, Amplex Red, an H_2_O_2_-sensitive dye was used. Solutions of 50 µM Amplex Red, 1 U/mL Horseradish peroxidase, increasing dilutions of imaging buffer, and 30 μM H_2_O_2_ were made in a 96-well plate. Samples were incubated for 30 min in the dark. Samples were then excited at 530 nm using a fluorescence spectrophotometer (Molecular Devices, Spectramax Gemini XPS Microplate Reader) and fluorescence was measured at 590 nm emission.

To determine the absolute H_2_O_2_ concentration from absorbance measurements, samples were prepared with increasing imaging buffer dilutions (0.003-1x). The recovery fraction for each imaging buffer dilution was determined using the following equation:

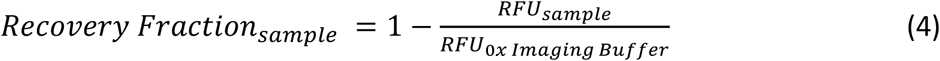

To predict the remaining H_2_O_2_ concentration in the 0, 0.5, and 1 mM H_2_O_2_ microtubule dynamics experiments, the mean recovery fraction was multiplied by each starting H_2_O_2_ concentration.

## Acknowledgements

MKG is supported by a National Institutes of Health grant NIGMS GM-103833 of the National Institutes of Health, RRG is supported by the National Institute of Health Training Program in Muscle Research (T32AR007612). JMM is supported by National Institutes of Health grant NHLBI R01 HL123874. Parts of this work were carried out in the Characterization Facility, University of Minnesota, a member of the NSF-funded Materials Research Facilities Network (www.mrfn.org) via the MRSEC program. We thank members of the Gardner laboratory for helpful discussions, and Dr. Taylor Reid for software assistance and guidance. We thank Dr. Thomas Surrey for the generous gift of Mal3 constructs.

## Author Contributions

Conceptualization: R.R.G. and M.K.G.

Methodology: all authors

Software: C.H. and M.K.G.

Formal Analysis: R.R.G., M.M., K.W., C.H., and M.K.G

Investigation: R.R.G., M.M., and K.W

Resources: B.R.T, H.V., H.C, J.M.M.

Data Curation: M.K.G.

Writing – Original Draft: R.R.G

Writing – Review & Editing: all authors

Supervision: M.K.G. and J.M.

Funding Acquisition: M.K.G. and J.M.

## Declaration of Interests

The authors declare no competing interests.

**Supplemental Movie 1: *In vitro* Dynamic Microtubule Assay**

A dynamic microtubule in a cell-free assay, imaged via TIRF microscopy. Stabilized microtubule seeds (red) are immobilized to an imaging chamber via anti-rhodamine antibody. Free tubulin (green) was introduced into the chamber in the presence or absence of H_2_O_2_. Movies acquired at 0.2 frames/sec using TIRF microscopy (0.16 μm pixel size) were used to make kymographs in Fig. 2A.

## References

Akella, J.S., Wloga, D., Kim, J., Starostina, N.G., Lyons-Abbott, S., Morrissette, N.S., Dougan, S.T., Kipreos, E.T., and Gaertig, J. (2010). MEC-17 is an α-tubulin acetyltransferase. Nature 467, 218–222.

Aumeier, C., Schaedel, L., Gaillard, J., John, K., Blanchoin, L., and Théry, M. (2016). Self-repair promotes microtubule rescue. Nat. Cell Biol. 18, 1054–1064.

Bauer, E., Klövekorn, W.-P., Schlepper, M., Schaper, W., Schaper, J., Heling, A., Zimmermann, R., Kostin, S., Maeno, Y., Hein, S., et al. (2000). Human Myocardium Increased Expression of Cytoskeletal, Linkage, and Extracellular Proteins in Failing Increased Expression of Cytoskeletal, Linkage, and Extracellular Proteins in Failing Human Myocardium. Circ Res 86, 846–853.

Castoldi, M., and Popov, A. V. (2003). Purification of brain tubulin through two cycles of polymerization-depolymerization in a high-molarity buffer. Protein Expr. Purif. 32, 83–88.

Chen, C.Y., Caporizzo, M.A., Bedi, K., Vite, A., Bogush, A.I., Robison, P., Heffler, J.G., Salomon, A.K., Kelly, N.A., Babu, A., et al. (2018). Suppression of detyrosinated microtubules improves cardiomyocyte function in human heart failure. Nat. Med. 24, 1225–1233.

Clark, H.M., Hagedorn, T.D., and Landino, L.M. (2014). Hypothiocyanous acid oxidation of tubulin cysteines inhibits microtubule polymerization. Arch. Biochem. Biophys. 541, 67–73.

Coombes, C., Yamamoto, A., McClellan, M., Reid, T.A., Plooster, M., Luxton, G.W.G., Alper, J., Howard, J., and Gardner, M.K. (2016). Mechanism of microtubule lumen entry for the α-tubulin acetyltransferase enzyme αTAT1. Proc. Natl. Acad. Sci. U. S. A. 113, E7176–E7184.

Coombes, C.E., Yamamoto, A., Kenzie, M.R., Odde, D.J., and Gardner, M.K. (2013). Evolving tip structures can explain age-dependent microtubule catastrophe. Curr. Biol. 23, 1342–1348.

Cooper, G. (2006). Cytoskeletal networks and the regulation of cardiac contractility: microtubules, hypertrophy, and cardiac dysfunction. Am. J. Physiol. Hear. Circ. Physiol. 291, H1003–14.

Desai, A., and Mitchison, T.J. (1997). Microtubule polymerization dynamics. Annu Rev. Cell Dev. Biol. 13, 83–117.

Dimitrov, A., Quesnoit, M., Moutel, S., Cantaloube, I., Poüs, C., and Perez, F. (2008). Detection of GTP-tubulin conformation in vivo reveals a role for GTP remnants in microtubule rescues. Science 322, 1353–1356.

Drum, B.M.L., Yuan, C., Li, L., Liu, Q., Wordeman, L., and Santana, L.F. (2016). Oxidative stress decreases microtubule growth and stability in ventricular myocytes. J. Mol. Cell. Cardiol. 93, 32–43.

Dye, R.B., Flicker, P.F., Lien, D.Y., and Williams, R.C. (1992). End-stabilized microtubules observed in vitro: Stability, subunit interchange, and breakage. Cell Motil. Cytoskeleton 21, 171–186.

Eshun-Wilson, L., Zhang, R., Portran, D., Nachury, M. V, Toso, D.B., Löhr, T., Vendruscolo, M., Bonomi, M., Fraser, J.S., and Nogales, E. (2019). Effects of α-tubulin acetylation on microtubule structure and stability. Proc. Natl. Acad. Sci. U. S. A. 116, 10366–10371.

Fernández-Puente, E., Sánchez-Martín, M.A., de Andrés, J., Rodríguez-Izquierdo, L., Méndez, L., and Palomero, J. (2020). Expression and functional analysis of the hydrogen peroxide biosensors HyPer and HyPer2 in C2C12 myoblasts/myotubes and single skeletal muscle fibres. Sci. Rep. 10, 871.

de Forges, H., Pilon, A., Cantaloube, I., Pallandre, A., Haghiri-Gosnet, A.M., Perez, F., and Poüs, C. (2016). Localized Mechanical Stress Promotes Microtubule Rescue. Curr. Biol. 26, 3399–3406.

Gardner, M.K. (2016). Cell Biology: Microtubule Collisions to the Rescue. Curr. Biol. 26, R1287–R1289.

Gardner, M.K., Charlebois, B.D., J??nosi, I.M., Howard, J., Hunt, A.J., and Odde, D.J. (2011a). Rapid microtubule self-assembly kinetics. Cell 146, 582–592.

Gardner, M.K., Zanic, M., Gell, C., Bormuth, V., and Howard, J. (2011b). Depolymerizing kinesins Kip3 and MCAK shape cellular microtubule architecture by differential control of catastrophe. Cell 147, 1092–1103.

Gell, C., Bormuth, V., Brouhard, G.J., Cohen, D.N., Diez, S., Friel, C.T., Helenius, J., Nitzsche, B., Petzold, H., Ribbe, J., et al. (2010). Microtubule dynamics reconstituted in vitro and imaged by single-molecule fluorescence microscopy. Methods Cell Biol. 2010, 221–245.

Giorgio, M., Trinei, M., Migliaccio, E., and Pelicci, P.G. (2007). Hydrogen peroxide: a metabolic by-product or a common mediator of ageing signals? Nat. Rev. Mol. Cell Biol. 8, 722–728.

Le Grand, M., Rovini, A., Bourgarel-Rey, V., Honore, S., Bastonero, S., Braguer, D., and Carre, M. (2014). ROS-mediated EB1 phosphorylation through Akt/GSK3β pathway: implication in cancer cell response to microtubule-targeting agents. Oncotarget 5, 3408–3423.

Guo, H., Xu, C., Liu, C., Qu, E., Yuan, M., Li, Z., Cheng, B., and Zhang, D. (2006). Mechanism and dynamics of breakage of fluorescent microtubules. Biophys. J. 90, 2093–2098.

Hein, S., Kostin, S., Heling, A., Maeno, Y., and Schaper, J. (2000). The role of the cytoskeleton in heart failure. Cardiovasc. Res. 45, 273–278.

Hinshaw, D.B., Miller, M.T., Omann, G.M., Beals, T.F., and Hyslop, P.A. (1993). A cellular model of oxidant-mediated neuronal injury. Brain Res. 615, 13–26.

Hori, M., and Nishida, K. (2009). Oxidative stress and left ventricular remodelling after myocardial infarction. Cardiovasc. Res. 81, 457–464.

Howard, J., and Hyman, A.A. (2003). Dynamics and mechanics of the microtubule plus end. Nature 422, 753–758.

Howarth, F.C., Calaghan, S.C., Boyett, M.R., and White, E. (1999). Effect of the microtubule polymerizing agent taxol on contraction, Ca2+ transient and L-type Ca2+ current in rat ventricular myocytes. J. Physiol. 516, 409–419.

Huang, B.K., and Sikes, H.D. (2014). Quantifying intracellular hydrogen peroxide perturbations in terms of concentration. Redox Biol. 2, 955–962.

Iwai, K., Hori, M., Kitabatake, A., Kurihara, H., Uchida, K., Inoue, M., and Kamada, T. (1990). Disruption of microtubules as an early sign of irreversible ischemic injury. Immunohistochemical study of in situ canine hearts. Circ Research. 67, 694–706.

Janke, C., and Montagnac, G. (2017). Causes and Consequences of Microtubule Acetylation. Curr. Biol. 27, R1287–R1292.

Kerr, J.P., Robison, P., Shi, G., Bogush, A.I., Kempema, A.M., Hexum, J.K., Becerra, N., Harki, D.A., Martin, S.S., Raiteri, R., et al. (2015). Detyrosinated microtubules modulate mechanotransduction in heart and skeletal muscle. Nat. Commun. 6, 1–10.

Kirschner, M., and Mitchison, T. (1986). Beyond self-assembly: From microtubules to morphogenesis. Cell 45, 329–342.

Kirschner, M., and Mitchison, T.J. (2002). Microtubule dynamics. J. Cell Sci. 115, 3–4.

Kormendi, V., Szyk, A., Piszczek, G., and Roll-Mecak, A. (2012). Crystal structures of tubulin acetyltransferase reveal a conserved catalytic core and the plasticity of the essential N terminus. J. Biol. Chem. 287, 41569–41575.

Kratzer, E., Tian, Y., Sarich, N., Wu, T., Meliton, A., Leff, A., and Birukova, A.A. (2012). Oxidative stress contributes to lung injury and barrier dysfunction via microtubule destabilization. Am. J. Respir. Cell Mol. Biol. 47, 688–697.

Landino, L.M., Hasan, R., McGaw, A., Cooley, S., Smith, A.W., Masselam, K., and Kim, G. (2002). Peroxynitrite oxidation of tubulin sulfhydryls inhibits microtubule polymerization. Arch. Biochem. Biophys. 398, 213–220.

Landino, L.M., Hagedorn, T.D., Kim, S.B., and Hogan, K.M. (2011). Inhibition of tubulin polymerization by hypochlorous acid and chloramines. Free Radic. Biol. Med. 50, 1000–1008.

Leibler, S., and Dogterom, M. (1993). Physical Aspects of the Growth and Regulation of Microtubule Structures. Phys. Rev. Lett. 70, 1347–1350.

Luduena, R.F., and Roach, M.C. (1991). Tubulin sulfhydryl groups as probes and targets for antimitotic and antimicrotubule agents. Pharmacol. Ther. 49, 133–152.

Ly, N., Elkhatib, N., Bresteau, E., Piétrement, O., Khaled, M., Magiera, M.M., Janke, C., Le Cam, E., Rutenberg, A.D., and Montagnac, G. (2016). αtAT1 controls longitudinal spreading of acetylation marks from open microtubules extremities. Sci. Rep. 6, 1–14.

Maruta, H., Greer, K., and Rosenbaum, J.L. (1986). The acetylation of alpha-tubulin and its relationship to the assembly and disassembly of microtubules. J. Cell Biol. 103, 571–579.

Maurer, S.P., Bieling, P., Cope, J., Hoenger, A., and Surrey, T. (2011). GTPγS microtubules mimic the growing microtubule end structure recognized by end-binding proteins (EBs). Proc. Natl. Acad. Sci. U. S. A. 108, 3988–3993.

Mellon, M.G., and Rebhun, L.I. (1976). Sulfhydryls and the in vitro polymerization of tubulin. J. Cell Biol. 70, 226–238.

Mitchison, T., and Kirschner, M. (1984). Dynamic instability of microtubule growth. Nature 312, 237–242.

Nishimura, S., Nagai, S., Katoh, M., Yamashita, H., Saeki, Y., Okada, J.I., Hisada, T., Nagai, R., and Sugiura, S. (2006). Microtubules modulate the stiffness of cardiomyocytes against shear stress. Circ. Res. 98, 81–87.

Nogales, E., Wolf, S.G., and Downing, K.H. (1998). Erratum: Structure of the αβ tubulin dimer by electron crystallography (correction) (Nature (1998) 391 (199-203)). Nature 393, 191.

Ohi, R., and Zanic, M. (2016). Ahead of the Curve: New Insights into Microtubule Dynamics. F1000Research 5, 314.

Poole, L.B., Klomsiri, C., Knaggs, S.A., Furdui, C.M., Nelson, K.J., Thomas, M.J., Fetrow, J.S., Daniel, L.W., and King, S.B. (2007). Fluorescent and affinity-based tools to detect cysteine sulfenic acid formation in proteins. Bioconjug. Chem. 18, 2004–2017.

Portran, D., Schaedel, L., Xu, Z., Théry, M., and Nachury, M. V (2017). Tubulin acetylation protects long-lived microtubules against mechanical ageing. Nat. Cell Biol. 19, 391–398.

Prins, K.W., Asp, M.L., Zhang, H., Wang, W., and Metzger, J.M. (2016). Microtubule-Mediated Misregulation of Junctophilin-2 Underlies T-Tubule Disruptions and Calcium Mishandling in mdx Mice. JACC Basic to Transl. Sci. 1, 122–130.

Reid, T.A., Schuster, B.M., Mann, B.J., Balchand, S.K., Plooster, M., McClellan, M., Coombes, C.E., Wadsworth, P., and Gardner, M.K. (2016). Suppression of microtubule assembly kinetics by the mitotic protein TPX2. J. Cell Sci. 129, 1319–1328.

Reid, T.A., Coombes, C., and Gardner, M.K. (2017). Manipulation and quantification of microtubule lattice integrity. Biol. Open 6, 1245–1256.

Reid, T.A., Coombes, C., Mukherjee, S., Goldblum, R.R., White, K., Parmar, S., McClellan, M., Zanic, M., Courtemanche, N., and Gardner, M.K. (2019). Structural state recognition facilitates tip tracking of eb1 at growing microtubule ends. Elife 8, 1–32.

Robison, P., and Prosser, B.L. (2017). Microtubule mechanics in the working myocyte. J. Physiol. 595, 3931–3937.

Robison, P., Caporizzo, M.A., Ahmadzadeh, H., Bogush, A.I., Chen, C.Y., Margulies, K.B., Shenoy, V.B., and Prosser, B.L. (2016). Detyrosinated microtubules buckle and bear load in contracting cardiomyocytes. Science (80-.). 352, aaf0659–aaf0659.

Sato, H., Nagai, T., Kuppuswamy, D., Narishige, T., Koide, M., Menick, D.R., and Cooper IV, G. (1997). Microtubule stabilization in pressure overload cardiac hypertrophy. J. Cell Biol. 139, 963–973.

Schaedel, L., John, K., Gaillard, J., Nachury, M. V, Blanchoin, L., and Théry, M. (2015). Microtubules self-repair in response to mechanical stress. Nat. Mater. 14, 1156–1163.

Schaedel, L., Triclin, S., Chrétien, D., Abrieu, A., Aumeier, C., Gaillard, J., Blanchoin, L., Théry, M., and John, K. (2019). Lattice defects induce microtubule self-renewal. Nat. Phys. 15, 830–838.

Schlieve, C.R., Lieven, C.J., and Levin, L.A. (2006). Biochemical activity of reactive oxygen species scavengers do not predict retinal ganglion cell survival. Investig. Ophthalmol. Vis. Sci. 47, 3878–3886.

Shida, T., Cueva, J.G., Xu, Z., Goodman, M.B., and Nachury, M. V (2010). The major α-tubulin K40 acetyltransferase αTAT1 promotes rapid ciliogenesis and efficient mechanosensation. Proc. Natl. Acad. Sci. U. S. A. 107, 21517–21522.

Sies, H. (2017). Hydrogen peroxide as a central redox signaling molecule in physiological oxidative stress: Oxidative eustress. Redox Biol. 11, 613–619.

Slezak, J., Tribulova, N., Pristacova, J., Uhrik, B., Thomas, T., Khaper, N., Kaul, N., and Singal, P.K. (1995). Hydrogen peroxide changes in ischemic and reperfused heart: Cytochemistry and biochemical and X-ray microanalysis Amer. J. Pathol. 147, 772–781.

Smyth, J.W., Hong, T.T., Gao, D., Vogan, J.M., Jensen, B.C., Fong, T.S., Simpson, P.C., Stainier, D.Y.R., Chi, N.C., and Shaw, R.M. (2010). Limited forward trafficking of connexin 43 reduces cell-cell coupling in stressed human and mouse myocardium. J. Clin. Invest. 120, 266–279.

Tagawa, H., Koide, M., Sato, H., Zile, M.R., Carabello, B.A., and Cooper, G. (1998). Cytoskeletal Role in the Transition From Compensated to Decompensated Hypertrophy During Adult Canine Left Ventricular Pressure Overloading. Circ. Res. 82, 751–761.

Thompson, J.A., and Hess, M.L. (1986). The oxygen free radical system: A fundamental mechanism in the production of myocardial necrosis. Prog. Cardiovasc. Dis. 28, 449–462.

Thompson, B.R., Martindale, J., and Metzger, J.M. (2016). Sarcomere neutralization in inherited cardiomyopathy: small-molecule proof-of-concept to correct hyper-Ca 2-sensitive myofilaments. yAm J Physiol Hear. Circ Physiol 311, 36–43.

Thompson, B.R., Soller, K.J., Vetter, A., Yang, J., Veglia, G., Bowser, M.T., and Metzger, J.M. (2019). Cytoplasmic nucleic acid-based XNAs directly enhance live cardiac cell function by a Ca 2+ cycling-independent mechanism via the sarcomere. J. Mol. Cell. Cardiol. 130, 1–9.

Tropini, C., Roth, E.A., Zanic, M., Gardner, M.K., and Howard, J. (2012). Islands containing slowly hydrolyzable GTP analogs promote microtubule rescues. PLoS One 7, e30103.

Tsutsui, H., Ishihara, K., and Cooper IV, G. (1993). Cytoskeletal role in the contractile dysfunction of hypertrophied myocardium. Science. 260, 682–687.

Tsutsui, H., Tagawa, H., Kent, R.L., McCollam, P.L., Ishihara, K., Nagatsu, M., and Cooper IV, G. (1994). Role of microtubules in contractile dysfunction of hypertrophied cardiocytes. Circ. 90, 533–555.

Valen, G., Sondén, A., Vaage, J., Malm, E., and Kjellström, B.T. (1999). Hydrogen peroxide induces endothelial cell atypia and cytoskeleton depolymerization. Free Radic. Biol. Med. 26, 1480–1488.

Vemu, A., Szczesna, E., Zehr, E.A., Spector, J.O., Grigorieff, N., Deaconescu, A.M., and Roll-Mecak, A. (2018). Severing enzymes amplify microtubule arrays through lattice GTP-tubulin incorporation. Science (80-.). 361.

Wang, L., Lopaschuk, G.D., and Clanachan, A.S. (2008). H 2 O 2 -induced left ventricular dysfunction in isolated working rat hearts is independent of calcium accumulation. J. Mol. Cell. Cardiol. 45, 787–795.

Webster, D. (2002). Microtubules in Cardiac Toxicity and Disease. Cardiovasc. Toxicol. 2, 75–89.

Wei-Guo, H.U., and Qi-Ping, L.U. (2014). Impact of oxidative stress on the cytoskeleton of pancreatic epithelial cells. Exp. Ther. Med. 8, 1438–1442.

White, E. (2011). Mechanical modulation of cardiac microtubules. Pflügers Arch. Eur. J. Physiol. 462, 177–184.

Xie, J., Zhou, X., Hu, X., and Jiang, H. (2014). H2O2 evokes injury of cardiomyocytes through upregulating HMGB1. Hell. J. Cardiol. 55, 101–106.

Xu, Z., Schaedel, L., Portran, D., Aguilar, A., Gaillard, J., Peter Marinkovich, M., Théry, M., and Nachury, M. V (2017). Microtubules acquire resistance from mechanical breakage through intralumenal acetylation. Science. 356, 328–332.

Zadeh, A.D., Cheng, Y., Xu, H., Wong, N., Wang, Z., Goonasekara, C., Steele, D.F., and Fedida, D. (2009). Kif5b is an essential forward trafficking motor for the Kv1.5 cardiac potassium channel. J. Physiol. 587, 4565–4574.

Zanic, M., Stear, J.H., Hyman, A.A., and Howard, J. (2009). EB1 recognizes the nucleotide state of tubulin in the microtubule lattice. PLoS One 4, 7585.

Zanic, M., Widlund, P.O., Hyman, A.A., and Howard, J. (2013). Synergy between XMAP215 and EB1 increases microtubule growth rates to physiological levels. Nat. Cell Biol. 15, 688–693.

Zhang, Y., Liu, X., Zhang, L., Li, X., Zhou, Z., Jiao, L., Shao, Y., Li, M., Leng, B., Zhou, Y., et al. (2018). Metformin Protects against H2O2-Induced Cardiomyocyte Injury by Inhibiting the miR-1a-3p/GRP94 Pathway. Mol. Ther. – Nucleic Acids 13, 189–197.

Zhao, J., Gao, J.L., Zhu, J.X., Zhu, H. Bin, Peng, X., Jiang, M., Fu, Y., Xu, J., Mao, X.H., Hu, N., et al. (2019). The different response of cardiomyocytes and cardiac fibroblasts to mitochondria inhibition and the underlying role of STAT3. Basic Res. Cardiol. 114, 1–16.

Zhou, C., Yan, L., Zhang, W. hui, and Liu, Z. (2019). Structural basis of tubulin detyrosination by VASH2/SVBP heterodimer. Nat. Commun. 10, 1–8.

